# Atypical Replication Error Pattern and Limited Repair Efficiency Contribute to Elevated Mutation Rate in Phage lambda

**DOI:** 10.1101/2025.08.29.673000

**Authors:** Julien Lopez, Magali Ventroux, Valentin Loux, Mahendra Mariadassou, François Lecointe, Rafael Sanjuan, Marina Elez, Marianne De Paepe

## Abstract

Understanding the rate and nature of spontaneous mutations is crucial for understanding and modulating the pace and trajectory of evolution. Yet, both remain poorly characterized in double-stranded DNA (dsDNA) phages, despite their relevance for phage-based therapies. Here, we address this gap for the dsDNA phage lambda, using four complementary approaches: mutation accumulation assay combined with whole-genome sequencing, mutation visualization assay, duplex sequencing and fluctuation assay. We find that the mutation rate of wild-type phage lambda is 4.9 ± 1.8 x10^-9^ per base per replication, approximately 15 times lower than previously estimated and about 20 times higher than that of its host, *Escherichia coli* (*E. coli*). Inactivation of Mismatch Repair (MMR), a major conserved cellular system for mutation avoidance, increases the lambda mutation rate by only 2-to 10-fold, in contrast to the approximately 150-fold increase in *E. coli*, however lambda does not exhibit the characteristic mutational bias associated with MMR deficiency. Interestingly replication of the lambda genome generates an error spectrum distinct from that of *E. coli*, characterized by a marked increase in transversions, poorly repaired by MMR. Together, these results reveal that lambda exhibits a replication error profile that is less amenable to repair, likely contributing to its elevated mutation rate.

## Introduction

Viruses evolve much faster than their hosts due to significantly higher mutation and recombination rates (Sanjuán et al. 2010). While host organisms typically exhibit mutation rates between 10⁻^9^ and 10⁻^10^ per base per replication (Lynch 2010), viral mutation rates are substantially higher, ranging from 10⁻³ to 10⁻⁸ (Duffy et al. 2008; Sanjuán and Domingo-Calap 2016). In living cells, spontaneous mutations are largely controlled by three key systems: (i) DNA polymerase proofreading activity, (ii) the Mismatch Repair (MMR) system, which corrects most residual polymerase errors, and (iii) repair pathways targeting oxidative DNA damage, such as MutTMY (Lee et al. 2012; Foster et al. 2015; Niccum et al. 2018; Zou et al. 2021). The high mutation rates of RNA viruses are largely attributed to their error-prone RNA-dependent polymerases, which lack proofreading activity, and the absence of RNA repair pathways (Duffy et al. 2008). Surprisingly, however, dsDNA phages, which rely on high-fidelity DNA polymerases for replication, also exhibit elevated mutation rates, and the underlying mechanisms remain unclear. Obtaining accurate estimates of mutation rates and spectra is a critical first step toward understanding these mechanisms, yet such data are currently lacking.

Lambda phage is one of the most extensively studied dsDNA phages. As a foundational model in molecular biology, its study has greatly advanced our understanding of gene regulation and recombination. Despite its scientific importance, a single estimate of its mutation rate and spectrum have been reported, both obtained through a genetic approach (Drake 1991; Wagner and Nohmi 2000). Specifically, mutation rate was estimated via a Fluctuation Assay (FA) and mutation spectrum through a selection of CII inactivation mutants followed by Sanger sequencing of *cII* gene. Mutation rate estimates derived from FAs, rely on selection to detect mutants, arising within a small genomic region (< 3 kb) (Luria and Delbrück 1943), and use of statistical models to infer the number of mutational events from the observed number of mutants (Foster 2006; Zheng 2015). These models rely on several assumptions, such as exponential growth without cell death, immediate expression of mutations, and detection of all mutants (Foster 2006), which might not all hold under all experimental conditions thereby leading to potentially inaccurate mutation rate estimate. The phenotypic mutation rate can be converted into a per-base mutation rate by estimating the target size, the number of distinct base changes that produce the phenotype, typically via sequencing hundreds of independent mutants (Drake et al. 1998; Lang and Murray 2008). However, the precision of such estimates is constrained by regional differences in mutation rates observed across the genome (Foster et al. 2013; Long et al. 2014; Dillon et al. 2015, 2017, 2018).

Lambda and *E. coli* share the same replicative DNA polymerase, Pol III (Zylicz et al. 1989), implying a potentially similar intrinsic replication fidelity but so far this hasn’t been investigated experimentally. Moreover, whether the host’s DNA repair systems, responsible for preserving genome integrity, function with comparable efficiency on the phage genome remains unknown. In the case of MMR, a key determinant of efficiency is the ability to distinguish the newly synthesized DNA strand from the template strand (Dohet et al. 1986). In *E. coli*, this discrimination relies on the transient undermethylation of GATC sites on the nascent strand, as Dam methylase trails behind Pol III during replication (Dreiseikelmann et al. 1979; Campbell and Kleckner 1988). In the absence of GATC methylation, this distinction is lost, rendering both DNA strands susceptible to cleavage by MutH (Au et al. 1992). Notably, previous studies have shown that lambda DNA is undermethylated (Hattman 1972; Szyf et al. 1984), and that GATC sites are underrepresented (Hénaut et al. 1996); which may impair the MMR ability to distinguish strands, thereby reducing error correction and increasing the mutation rate.

Here, we investigated mutagenesis in lambda phage using four complementary approaches: mutation accumulation coupled with whole-genome sequencing (MA-WGS), as assay to visualize replication error in infected cells, a sequencing method for detecting ultra-rare mutations, Duplex Sequencing (DS), and a fluctuation assay (FA). Together, this enabled us to i) obtain an accurate estimate of lambda’s mutation rate, ii) characterize its mutation spectrum and iii) uncover the replication errors pattern on lambda genome and characterize its repair.

## Results

### Reassessment of lambda mutation rate using genomic methods

To investigate mutagenesis of lambda, we employed MA+WGS, the state-of-the-art approach widely used to estimate mutation rates in diverse organisms (Lynch et al. 2016), but not yet applied to phages. MA-WGS detects only fixed mutations, present in every individual of the population. As in other systems with low per-genome, per-replication mutation rates, applying MA-WGS on lambda requires extended propagation to allow sufficient mutations to arise and reach fixation. To limit natural selection, especially the loss of deleterious mutations, regular population bottlenecks are applied by initiating each new passage from a single individual. We established thirty MA lines of lambda by propagation on *E. coli* lawns grown on LB agar plates, streaking a single lysis plaque from each plate onto a fresh lawn for 150 passages (Table 1). The number of genome replications per line was estimated by regularly measuring plaque titers (approximately every 9 passages, Figure S1) and assuming exponential growth. While this model simplifies phage replication given that i) lambda does not replicate exclusively via bidirectional replication (Taylor and Wȩgrzyn 1995), and ii) not every phage successfully infects a cell, we estimate that these departure do not substantially bias mutation rate estimates (Supplementary Information 1.5).

**Table 1:** Estimates of lambda mutation rate from genomic analysis. Values in parenthesis indicate 95% confidence intervals (see Methods and Table S1 for calculation details). N, number; BPS, base pair substitutions

| Host genotype | Method | N lines | N DNA doublings | N BPS/line | N indels/line | Substitution rate (/base/ replication) | Mutation rate (/base/ replication) |
| --- | --- | --- | --- | --- | --- | --- | --- |
| WT | MA-WGS | 30 | 3,876 | 1.47 | 0.27 | $7.8 \times 10^{-9}$ | $9.2 \times 10^{-9}$ |
| WT | DS | 4 | 39 | 47 | - | $1.23 [1.03; 1.43] \times 10^{-8}$ | - |
| *mutS* | DS | 4 | 39 | 91 | - | $2.50 [2.20; 2.80] \times 10^{-8}$ | - |

We also estimated the mutation rate of lambda using DS, a highly accurate next-generation sequencing (NGS) approach that distinguishes true mutations from sequencing errors by comparing both DNA strands (Schmitt et al. 2012; Risso-Ballester et al. 2016). The ultra-low error rate (down to 1 x10^-8^) allows detection of rare mutations within a population, without the need for prolonged propagation and mutation fixation. Because it captures mutations shortly after they arise, DS introduces minimal selection bias and can even detect lethal mutations, as >90% of replications occur during the final lytic cycle. To apply DS to lambda, we generated four high-titer populations of lambda (∼5 x10^11^ phages each) by amplification from a single phage, and then extracted and sequenced the phage DNA. Per base pair substitution frequency in lambda ranged from 6.0 x10^-7^ to 8.1 x10^-7^, significantly higher than in the control sample (pUC plasmid) where no mutations were expected due to high replication fidelity (see Methods). This difference suggests that ∼80% of the mutations detected in lambda are genuine (Table S1), allowing estimation of the lambda mutation rate.

### Selection is minimal during MA and DS experiments with lambda

Reliable estimation of mutation rate using WGS relies on an unbiased sampling of spontaneous mutations. Of the 52 mutations detected in our MA-WGS experiment, only one was found in more than one lineage (present in two), suggesting the lack of strong adaptive selection. Consistently, the number of PFU per plaque remained stable over time (Figure S1), further supporting the lack of adaptive mutations in MA lines.

We analyzed the genomic distribution of mutations in both DS and MA-WGS datasets. Notably, 12 of the 30 MA lines exhibited clusters of mismatches in regions homologous to defective prophages DLP12, in 11 lines, and Qin, in 1 line. These mismatches precisely matched the corresponding prophage sequences, indicating homologous recombination events rather than mutation hotspots. As such, they were excluded from the calculation of the BPS rate. From the observed number of recombination events, we estimated the rate of horizontal gene transfer via homologous recombination between lambda and the Dlp12 prophage to be at least 3 x10^-3^ per replication, consistent with previous estimate based on a gene marker (De Paepe et al. 2014).

To assess whether mutations were uniformly distributed across the genomes, we calculated BPS frequencies for the DS dataset using the number of mutations in a 1 kb sliding window and compared local variations to the genome-wide average (Figure 1a). Per-window mutation rates appeared constant across the genome (Figure 1a). We also compared the number of mutations per transcription unit to expectations based on length of each unit and constant mutation rate (Figure S2a) and found no significant deviations. In addition, for both DS and MA-WGS datasets the distances between consecutive mutations exhibit no deviation from a Poisson distribution, again consistent with a uniform mutation rate across the genome (Figure 1b). Together, these results demonstrate the absence of mutational hotspots and selection in MA-WGS and DS experiments.

**Figure 1.**
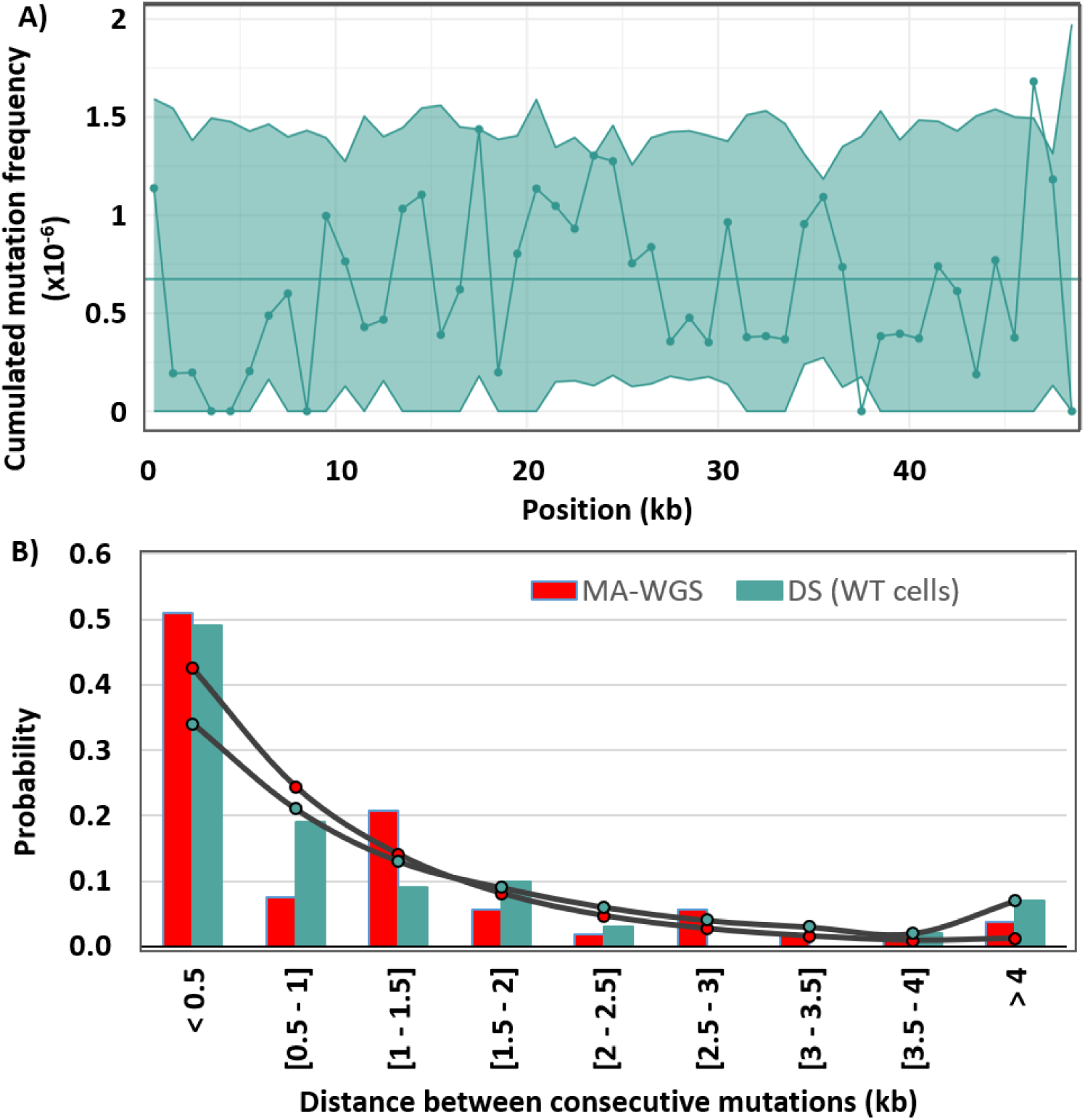
Uniform mutation frequency indicates absence of selection in MA-WGS and DS experiments on lambda. **A)** Mutation frequency across the lambda genome, estimated from the DS experiment. The horizontal line indicates the genome-wide estimated; where the dotline indicates mutation frequencies estimated using a 1 kb sliding window. Shaded area indicates 95% confidence interval, based on the the genome-wide estimate, for each window. **B)** Distribution of distances (discretized in bins of length 0.5 kb) between consecutive mutations from MA-WGS and DS datasets (bar plot), compared to the theoretical Poisson distribution (dotted line).

### Genomic analysis show that lambda mutation rate was previously overestimated

As MA-WGS and DS results were consistent with random, unbiased mutagenesis (Figure 1 & S2), we used them to estimate the mutation rate of lambda. From the MA-WGS data, we calculated BPS, indel and overall mutation rate by dividing the total number of observed mutations (BPS, indels or both) by the estimated number of genome replications across all MA lines. This yielded rates of 7.81 ± 0.09 x10^-9^; 1.42 ± 0.02 x10^-9^; 9.23 ± 0.11 x10^-9^ rate per base per replication for BPS, indels and overall mutations respectively (Table 1). As detailed in Supplementary Information 1.4 & 1.5, these represent upper-bound estimates, since the total number of replications is likely underestimated under the assumption of perfect exponential replication during phage growth.

For the DS dataset, analysis of mutation frequencies by type indicated that only BPS could be reliably estimated, as indel frequencies were too low to be distinguished from background noise (Table S1), consistent with findings in another dsDNA virus (Risso-Ballester et al. 2016). We therefore focused on BPS, calculating their frequency by subtracting the expected number of sequencing-error derived BPS from the total number of unique BPS, and dividing by the total number of bases sequenced. The BPS rate per replication was then obtained by dividing this frequency by the estimated number of phage genome replications, resulting in a rate of 1.23 x10^-8^ per replication (Table 1). As with the MA-WGS, this value is similarly considered an upper estimate, given the assumptions made on phage replication (Supplementary Information 1.4)

The DS experiment confirmed the BPS mutation rate estimated by MA-WGS, with the DS derived value falling within the expected range of methodological variation (50% higher). Together, these results indicate that the mutation rate of lambda is below 9.2 x10^-9^ per base per replication, approximately 8-fold lower than the previous estimate obtained using a fluctuation assay (Drake 1991).

### Fluctuation assay confirms lambda mutation rate is lower than previously estimated

Genomic approaches for estimating mutation rates are reliable, but either long and fastidious (MA+WGS) or expensive (DS), limiting their use across multiple conditions. In addition, in the context of lambda phage, MA-WGS yields only a small number of mutations within the manageable number of lines and experimental duration, while DS is prone to sequencing errors, both of which hinder accurate determination of the mutation spectrum, that is, the relative frequency of each mutation type. To overcome these limitations, we employed a FA, which is not subject to the aforementioned constraints. For this assay, we used the *cII* selection system of lambda (Jakubczak et al. 1996), where CII inactivation results in a selectable phenotype, the ability to form lysis plaques on an *E. coli* FtsH deficient host. We performed 10 fluctuation assays, each comprising 17 to 96 independent cultures. The distribution of mutant numbers across cultures was consistent with exponential replication, i.e., follows a Luria-Delbrück distribution (Figure S3). We therefore estimated the phenotypic mutation rate of CII inactivation per replication using the Ma–Sandri–Sarkar Maximum Likelihood Estimator (MSS-MLE) method. This model assumes a strictly bidirectional replication (genomic estimates mentioned above are also based on this assumption). For comparison, we also applied the historical *P_0_*method, which does not require assumptions about replication mode. The results were in close agreement (Table S2), supporting the robustness of our mutation rate estimates.

To convert the phenotypic mutation rate to a per-base mutation rate, we sequenced the *cII* gene in a large number of mutants to determine the target size, i.e., the number of nucleotides whose modification results in a null CII phenotype. Among 656 sequenced mutants, 425 harbored mutations in a 6G repeat, while only 138 were BPS, distributed along the gene. To improve the detection of BPS, we engineered a lambda strain in which the 6G repeat in *cII* gene was disrupted by replacing one G with an A, without altering the amino acid sequence. Sequencing the *cII* gene from 198 mutants of this phage revealed an additional 135 BPS. This expanded dataset provided sufficient coverage to apply the method of Lang & Murray (Lang and Murray 2008) for estimating the target size, and in turn, the per base mutation rate (Table S2). Briefly, we first identified the unique mutations responsible for the phenotype, categorized them by type (e.g. nonsense, missense, indels) and calculated the effective target size for each class, defined as the number of base positions where a mutation results in a detectable phenotype. A weighted average target size was then computed based on the relative contribution of each mutation type (e.g. nonsense + missense for BPS). Dividing the phenotypic mutation rate by this average target size yielded a per-base, per-replication BPS. The results were similar between the two lambda strains: 7.37 ± 0.91 x10^-9^ for lambda WT and 4.42 ± 1.61 x10^-9^ for lambda *cII*_G183A_ (Table 2). Due to the indel hotspot in the WT *cII* gene, the overall mutation rate (BPS + indels) was 4.4-fold higher in the WT *cII* than in the mutated version (Table 2).

**Table 2:** Estimate of lambda mutation rate from fluctuation assay. Values in parenthesis indicate 95% confidence intervals (see Methods for calculation details; full data are provided in Tables S2 & S3)

| lambda strain | Host strain | Phenotypic mutation rate per cycle ($\times 10^{-6}$) | Substitution rate ($\times 10^{-9}$ /base/ replication) | Mutation rate ($\times 10^{-9}$ /base/ replication) |
| --- | --- | --- | --- | --- |
| lambda WT | WT | 5.57 [4.88; 6.26] | 7.33 [6.45; 8.28] | 21.5 [18.9; 24.2] |
| lambda WT | *mutS* | 53.1 [37.94; 68.34] | 48.5 [34.60; 62.40] | 203.0 [145.0; 261.0] |
| lambda *cII*<sub>G183A</sub> | WT | 0.88 [0.56; 1.20] | 4.42 [2.81; 6.03] | 4.90 [3.12; 6.68] |
| lambda *cII*<sub>G183A</sub> | *mutS* | 9.07 [6.69; 11.45] | 45.5 [33.6; 57.5] | 50.5 [37.3; 63.8] |

The BPS rate obtained using the FA is 2-fold lower than the rate estimated by MA-WGS. This is consistent with the fact that MA-WGS-based rate represents an upper estimate (see Supplementary Information 1.4 & 1.5). Based on these results, we estimate that the mutation rate of lambda is about ∼5 x10^-9^ per base per replication, ∼16-fold lower than the previously reported estimate derived from *cI*-based fluctuation assay (Dove 1968; Drake 1991). For comparison, the mutation rate of *E. coli* has been estimated at 2.2 x10^-10^ mutations per base per generation per replication (1.88 x10^-10^± 0.46 and 2.45 x10^-10^ ± 0.49 for two *E. coli* K12 strains) using the MA-WGS approach (Lee et al. 2012). Our results indicate that lambda phage mutates ∼20 times faster than *E. coli*.

### Lambda mutation rate is minimally affected by the inactivation of MMR

We next investigated whether the elevated mutation rate of lambda could be attributed to inefficient MMR, due to undermethylation of its genome. This hypothesis leads to four key predictions: (i) inactivation of MMR should have a much weaker effect on the mutation rate of lambda compared to that of *E. coli*, (ii) mismatch detection by MutS should remain efficient, (iii) enhancing Dam-mediated methylation of the lambda genome should reduce its mutation rate, and (iv) the mutation spectrum of lambda should exhibit signatures of defective MMR.

To test the first prediction, we compared the mutation rate of lambda infecting WT *vs* MMR deficient *E. coli*, using DS (Table 1). Consistent with our hypothesis, MMR inactivation led to a modest 2-fold increase in *lambda*’s BPS mutation rate (Table 1). This contrasts sharply with the effect of MMR loss typically observed in bacteria, which often results in mutation rate increases exceeding 100-fold (Long et al. 2018). To validate this modest effect, we independently estimated lambda’s mutation rate using FA with the WT *cII* allele. Consistent with the DS results, the FA showed only a modest increase upon MMR inactivation, 6.6-fold for BPS and 9.4-fold for all mutations, using the WT phage, and 10.3-fold for BPS and all mutations, using the *cII*_G183A_ phage (Table 2), substantially lower than the ∼150 increase observed in *E. coli* (Lee et al. 2012; Niccum et al. 2018). These results indicate that lambda mutagenesis is only weakly affected by the host’s MMR activity.

### MMR detects replication errors on lambda genome

If the lambda genome is insufficiently methylated at GATC sites on the template strand, MMR is expected to fail at the repair step, while the recognition of mismatches, which primarily arise from replication errors during growth, should remain unaffected. To assess MMR’s ability to detect mismatches on the lambda genome, we employed a replication error visualization assay that we previously developed (Robert et al. 2018; Enrico Bena et al. 2024). This assay combines fluorescent labeling of the MMR protein MutL with live-cell time-lapse microscopy and a microfluidic device *mother machine* (Figure 2a), enabling replication errors to be visualized as bright spots of MutL, in individual *E. coli* cells under tightly controlled conditions.

**Figure 2.**
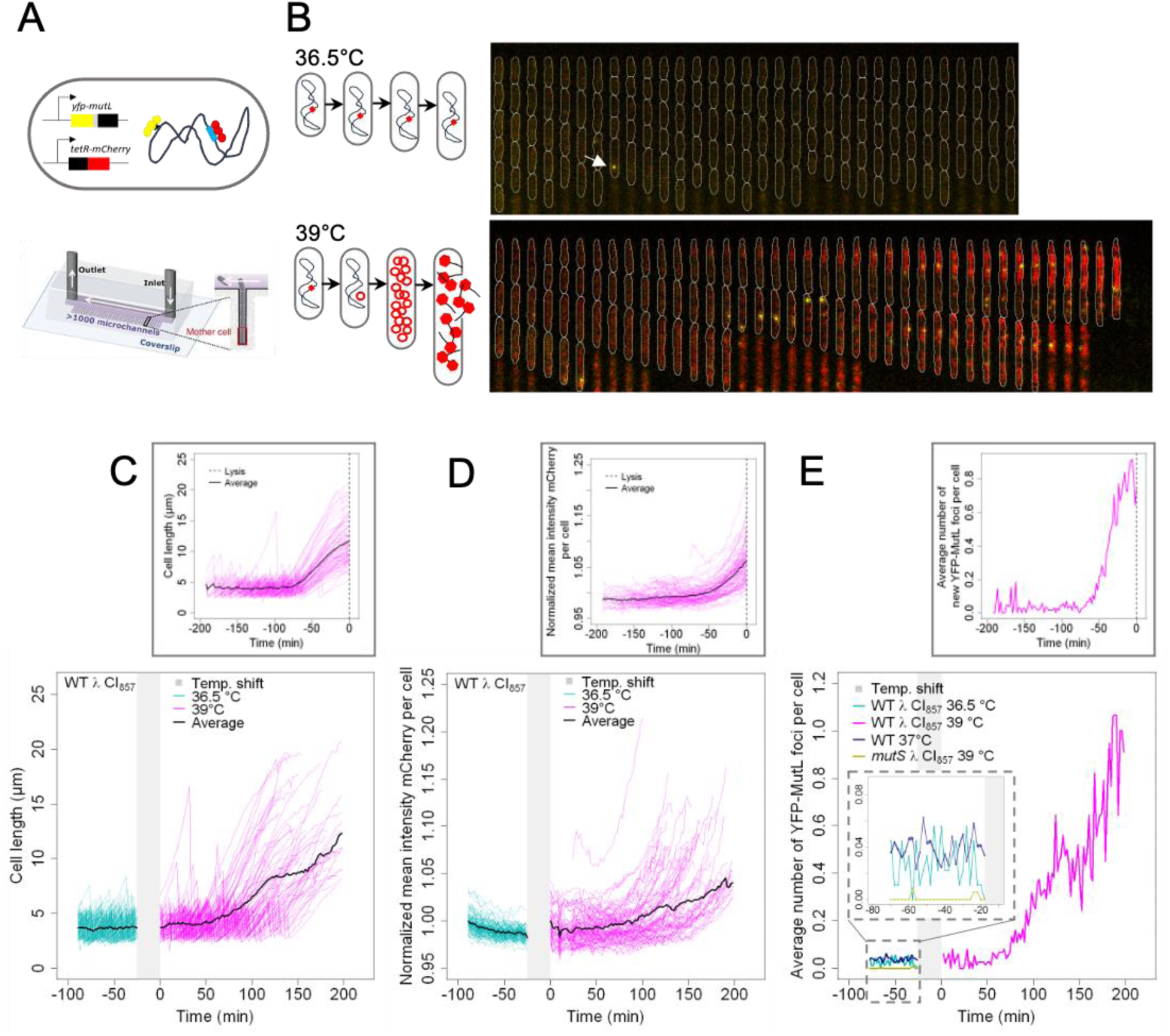
Visualization of replication errors during lambda infection. **A)** Schematic of the experimental setup. **Top**: *E. coli* strain expresses YFP-MutL from the chromosome and TetR-mCherry from a plasmid, and carries a lambda lysogen *cI_857_* containing 21 *tetO* arrays (blue). YFP-MutL (yellow) marks replication errors; TetR-mCherry (red) labels the *tetO* arrays. **Bottom**: Mother machine microfluidic device. **B) Left**: Schematics of lambda life cycles at 36.5 °C and 39 °C. **Right**: Corresponding kymographs. **Top**: At 36.5 °C, lambda remains in the lysogenic state (red square in the schematic); *E. coli* cells grow and divide, each retaining a single integrated lambda per genome. A YFP-MutL focus (white arrow) marks a replication error; cell contours are shown in white. **Bottom**: At 39 °C, the lytic cycle is rapidly induced: lambda genome excises, replicates, and accumulates (increased red fluorescence). The host cell elongates, fails to divide, and lyses (disappearance of cell), releasing mature virions. **C)** Cell length over time in a representative experiment using WT *E. coli*. **Bottom panel**: Each line represents one of 90 microchannels. Three experimental phases are highlighted: i) approximately 1 h at 36.5 °C (cyan), ii) a 25-min acquisition pause during temperature shift (shaded grey area), and iii) imaging resumed at 39 °C (magenta) for approximately 200 min until all cells lysed. Time 0 marks the start of the third phase. Solid black line shows the average cell length over time. **Top panel**: Same data from the 39 °C phase of the bottom panel, time-shifted so that all cells lysis events align at time 0, enhancing visualization of cell elongation prior to lysis. **D)** Same dataset as in C), showing mean mCherry fluorescence intensity per cell over time. Data are normalized to the value at the start of each acquisition phase. **Top panel** shows time-shifted data (as in C) to enhance visualization of fluorescence accumulation before lysis. **E)** Average number of newly appeared YFP-MutL foci per cell and per frame as a function of time. Cyan and magenta correspond to the same datasets shown in panels C) and D). The plot also includes data from two other representative experiments i) with WT *E. coli* (non-lysogen) grown at 37 °C (blue; data from Enrico Bena et al. 2024), showing a similar YFP-MutL foci rate before temperature shift in both lysogenic and non-lysogenic WT strain, ii) with *mutS E. coli* lysogen grown at 39 °C (yellow), showing that the YFP-MutL foci formed at this temperature are not non-functional aggregates, but rather mark replication errors on the DNA. **Top panel**: Same WT 39 °C dataset as in the bottom panel, but with time-shifting to align lysis events at time 0, highlighting the increase in replication errors before cell lysis. WT, wild-type *E. coli*; *mutS*, *E. coli* with inactivated *mutS*; λ cI857, lambda lysogen *cI_857_*.

We grew *E. coli* cells carrying a lambda prophage with the temperature-sensitive repressor *cI_857_* and expressing *yfp-mutL* from the Plac promoter in mother machine microchannels. In some experiments, we used a lysogenic strain containing 21 *tet0* binding sites inserted into the *bor* region of lambda (Trinh et al. 2020) and expressing constitutively TetR-mCherry from a plasmid (Figure 2b). This allowed fluorescent labeling of lambda DNA. To induce the lytic cycle, we increased the temperature from 36.5°C to 39°C (Bednarz et al. 2014) and monitored YFP-MutL foci together with tdCherry or mCherry fluorescence, before and after lambda induction (Figure 2b). Images were acquired every two minutes and analyzed using the BACMMAN software, enabling automated quantification of cells (via phase contrast or tdCherry), YFP-MutL foci and mCherry signal (Figure 2b).

Although the exact timing of prophage induction after the temperature shift and start of the lytic cycle could not be precisely determined, phage replication occurs during the last cell cycle, i.e., between the last cell division and cell lysis, as once lambda lytic cycle begins, cells cease dividing (Sergueev et al. 2002). During this phase, the initiation of new replication forks on the *E. coli* chromosome is blocked, although replication continues from pre-existing forks (Wold et al. 1982; Sergueev et al. 2002). Consistent with this, we observed a progressive increase in cell size during the final cell cycle (Figure 2c). In parallel, mCherry fluorescence, used as a proxy for lambda genome copy number, increased progressively during this phase, indicating active phage DNA replication (Figure 2d). Prior to the lytic cycle, the frequency of new YFP-MutL foci remained stable over time, consistent with earlier studies (Enrico Bena et al. 2024) (Figure 2e). In contrast, during the last cycle, we observed a marked increase in the frequency of YFP-MutL foci (Figure 2e), paralleling the increase in lambda genome copy number measured via by mCherry (Figure 2e) and quantitative PCR (Figure S4). This temporal concordance strongly suggests that the MMR system detects mismatches arising during lambda replication.

To assess the efficiency of MMR recognition of these mismatches on lambda, we compared the observed number of YFP-MutL foci per lytic cycle to the number of replication errors expected from lambda DNA synthesis. Since lambda is replicated by *E. coli* DNA polymerase III (Pol III), we estimated the number of replication errors based on the lambda genome size, the number of genomes replicated, and the per-base error rate of Pol III (Supplementary Information 1.1). As the intrinsic *in vivo* error rate of Pol III remains unknown, we used the mutation rate of MMR deficient *E. coli* cells as a proxy (3.26 ± 1.51 × 10^-8^ mutations per base, see Lee et al. 2012), as all replication errors are converted into mutations in this background. To compare this estimate with the number of YFP-MutL foci, which indicates number of replication errors rather than mutations, we adjusted for DNA segregation (Supplementary Information 1.2). We found that the number of YFP-MutL foci was approximately twice the number predicted based solely on the amount of lambda genome replication (Supplementary Information 1.1. & 1.2). This discrepancy may reflect contributions from concurrent *E. coli* genome replication (Supplementary Information 1.3) and/or higher error rate of Pol III during lambda lytic cycle. Based on our calculations, *E. coli* replication could at most account for less than half of the observed foci, assuming the Pol III error rate remains unchanged (Supplementary Information 1.1-1.3). Moreover, our data argue against a significant contribution from recombination-associated mismatches (Supplementary Information 1.3). Together, these findings suggest that a substantial number of replication errors on the lambda genome are effectively recognized by the MMR system.

### GATC undermethylation does not impair MMR in lambda

We next investigated whether increasing Dam-mediated methylation on the lambda genome could reduce its mutation rate. To test this, we multiplied lambda on three strains differing in Dam expression: a WT strain, a strain carrying an extra chromosomal copy of *dam* under the control of the PLtetO1 promoter, and a *dam* knockout strain used as a negative control (Figure 3). Mean adenine methylation at GATC sites on lambda genome increased from 46% in WT strain to 77 % in the Dam-overexpressing strain (Figure 3a). Methylation level across GATC sites was generally uniform across the lambda genome (Figure S5), except for a ∼5 kb region comprised between the integration site *attP* and the zone of convergence of the two main lambda promoters, P_L_ and P_R_, where methylation level dropped below 20% (Figure S5). Despite this local drop in methylation level, no corresponding increase in mutation frequency was observed in this region in either DS and MA experiments, indicating that regional variation in GATC methylation on lambda does not affect mutagenesis.

**Figure 3.**
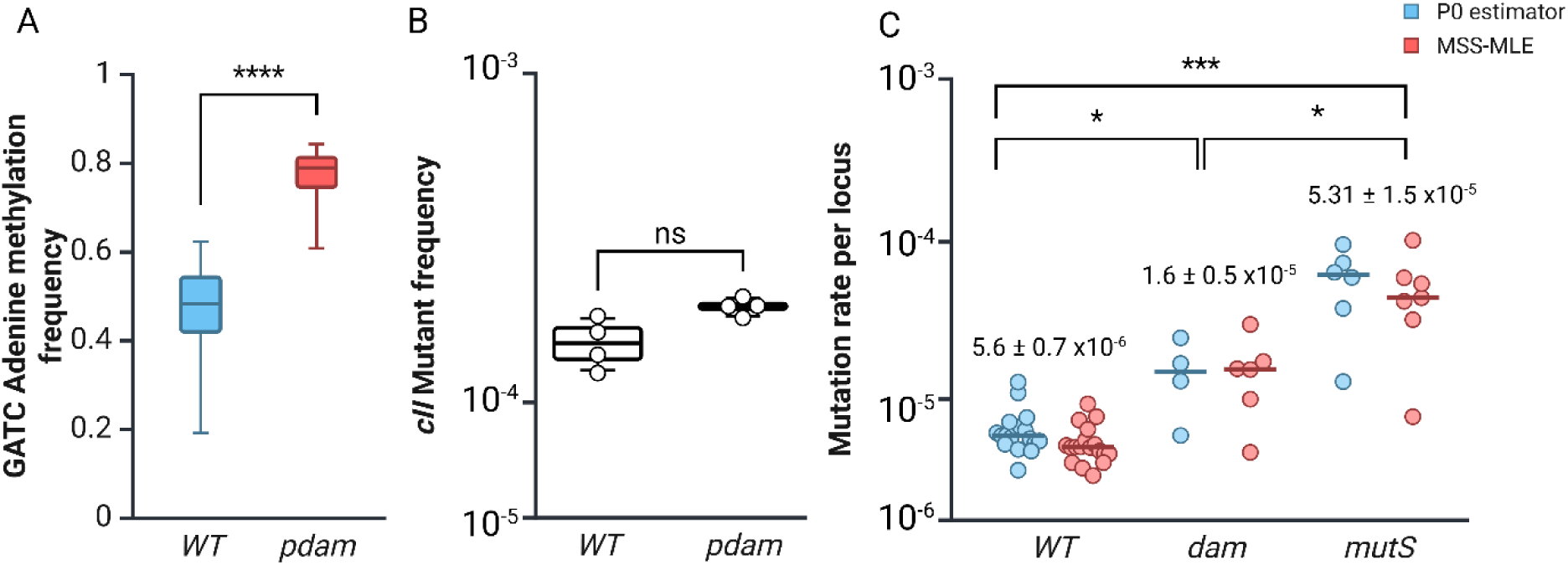
increased GATC methylation does not impact MMR efficiency in lambda. **A)** Boxplot showing the average percentage of adenine methylation at GATC sites in lambda. Lambda was propagated in WT cells or in cells overexpressing Dam (p*dam*). Two independent lysates were analysed per condition and pooled, as no significant differences were observed. **B)** Frequency of *cII* loss-of-function mutants in lambda lysates derived from WT or Dam overexpressing *E. coli*. Four independent lysates were analyzed per condition. **C)** Mutation rate at *cII* locus per replication, estimated using fluctuation assay and the P₀ method or MSS-MLE (Ma–Sandri–Sarkar Maximum Likelihood Estimator). Lambda was propagated on WT, *dam* deleted (*dam*) or *mutS* deleted (*mutS*) cells. Statistical comparisons were performed using the Mann-Whitney U-test (α = 0.05). ns = not significant.

To further assess whether increased methylation impacts mutation frequency, we performed *cII*-based FAs, comparing strains with or without Dam overexpression. Despite the higher methylation level, Dam overexpression did not result in a reduced mutant frequency (Figure 3b). In contrast, *dam* deletion led to a significant three-fold increase in *cII* mutant frequency (Figure 3c). This increase was notably smaller than the rise in *E. coli* mutagenesis reported following *dam* inactivation (Marinus and Morris 1975; Glickman and Radman 1980), consistent with our observation that MMR is less efficient on the lambda genome than on the *E. coli* chromosom*e* (Table 2). It was also lower than the increase observed when lambda replicated in MMR deficient *E. coli* cells (Figure 3c), indicating that, despite the ∼50% GATC methylation level, MMR still corrects a substantial fraction of replication errors in lambda. Therefore, 50% GATC methylation may be sufficient for strand discrimination by MMR, and other factors beyond GATC undermethylation likely contribute to the reduced MMR efficiency on lambda.

### The mutation spectrum of lambda reveals an atypical replication error pattern

If MMR inefficiency during the lambda lytic cycle were the primary cause of its elevated mutation rate, we would expect the lambda mutation spectrum to exhibit a characteristic signature of MMR deficiency. MMR is highly efficient at correcting replication errors that lead to transition mutations, commonly produced by replicative polymerases, but is less effective at repairing the rarer transversion errors (Long et al. 2018). In *E. coli*, the mutation spectrum of MMR deficient cells has been thoroughly characterized using both FA, using *lacI* and *rpoB* reporter genes (Leong et al. 1986; Schaaper and Dunn 1987), and MA-WGS (Lee et al. 2012; Niccum et al. 2018). Early studies of *lacI* mutations, which affect the lactose operon repressor, revealed a strong bias toward transitions, with 96% and 97% of mutations classified as transitions in datasets comprising 365 and 1,184 mutations respectively in (Schaaper and Dunn 1987) and (Leong et al. 1986). This strong transition bias was later confirmed by MA-WGS analyses, which reported 97% transitions in MMR deficient *E. coli* based on substantially larger mutation datasets (1,625 and 30,061 mutations respectively; Lee et al. 2012; Niccum et al. 2018).

To assess whether the lambda mutation spectrum exhibits features characteristic of MMR deficient *E. coli*, we analyzed mutations that arose during lambda replication. We excluded the DS dataset from the analysis due to its high error background. Instead, we focused on the 273 BPS identified in CII null mutants (Table 3) along with 44 BPS from our MA-WGS dataset (Table S4). In both cases, lambda replicating on WT *E. coli* showed a significantly lower proportion of transition mutations (<80%) compared to the strong transition bias reported in MMR deficient *E. coli* (Figure 4 & S6, Table 3, S3&4). Note that the proportion of transitions in an *E. coli* strain with partial MMR activity (*mutH* deficient cells expressing MutH from an inducible promoter showing a 78-fold increase in mutation rate relative to WT) closely matches that of fully MMR deficient strain (96% and 95% transitions respectively; Enrico Bena et al. 2024; Figure 4 & S6, Table S4).

**Figure 4.**
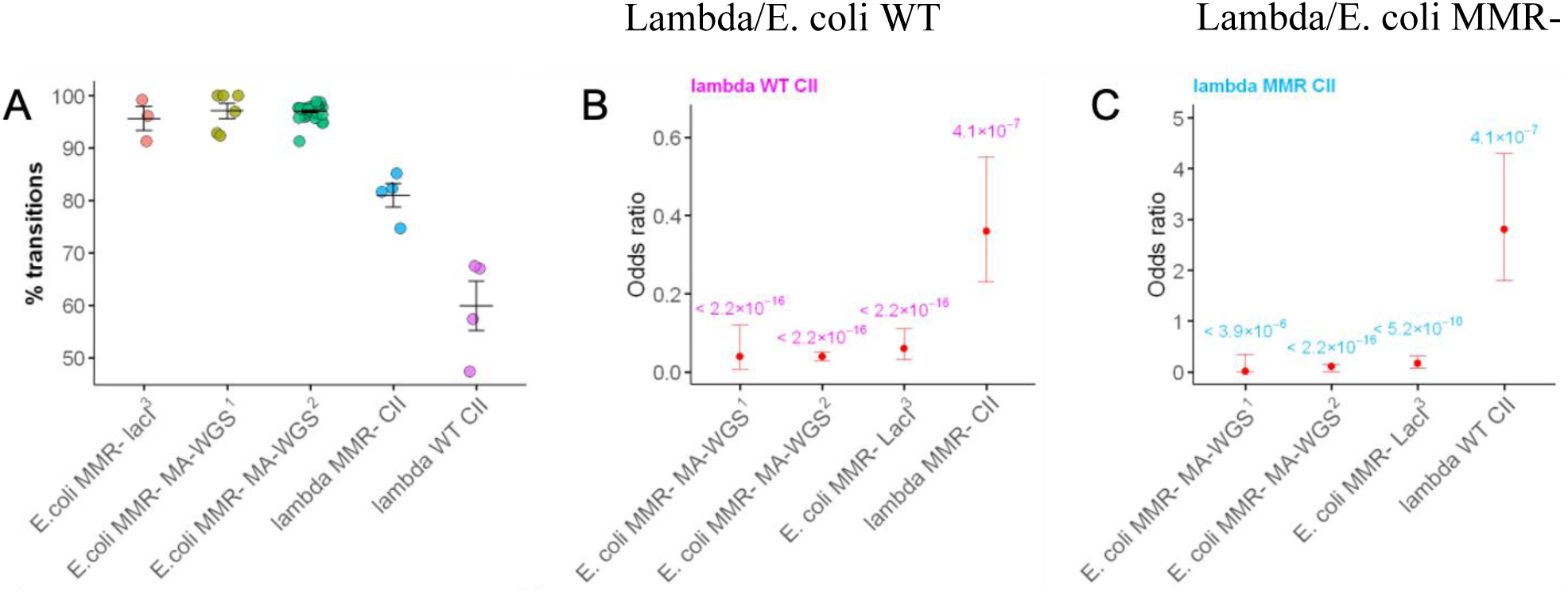
Lambda and E. coli chromosomes display a different mutation profile in MMR deficient strains. **A)** Percentage of transitions in lambda and MMR deficient *E. coli.* Mutations were obtained by selection for *cII* (lambda) or *lacI* (*E. coli*) inactivation or by MA-WGS (lambda and *E. coli*). Data for *E. coli* spectra are from: ^1^Enrico Bena et al., 2024, *Nature Communications*; ^2^Niccum et al., 2018, *Genetics*; ^3^Schaaper et al., 1987, *PNAS*. Each colored dot represents an independent experiment (≥3 replicates per group); black dots indicate group means; vertical lines indicate ± standard error of the mean. **B&C)** Statistical comparisons of transversion-to-transition ratios between lambda grown on WT (B) or MMR deficient (C) *E. coli* and the conditions indicated on the x-axis. Red dots indicate odds ratios from Fisher’s exact test; error bars represent 95% confidence intervals, p-values shown above dots indicate statistical significance.

**Table 3:**
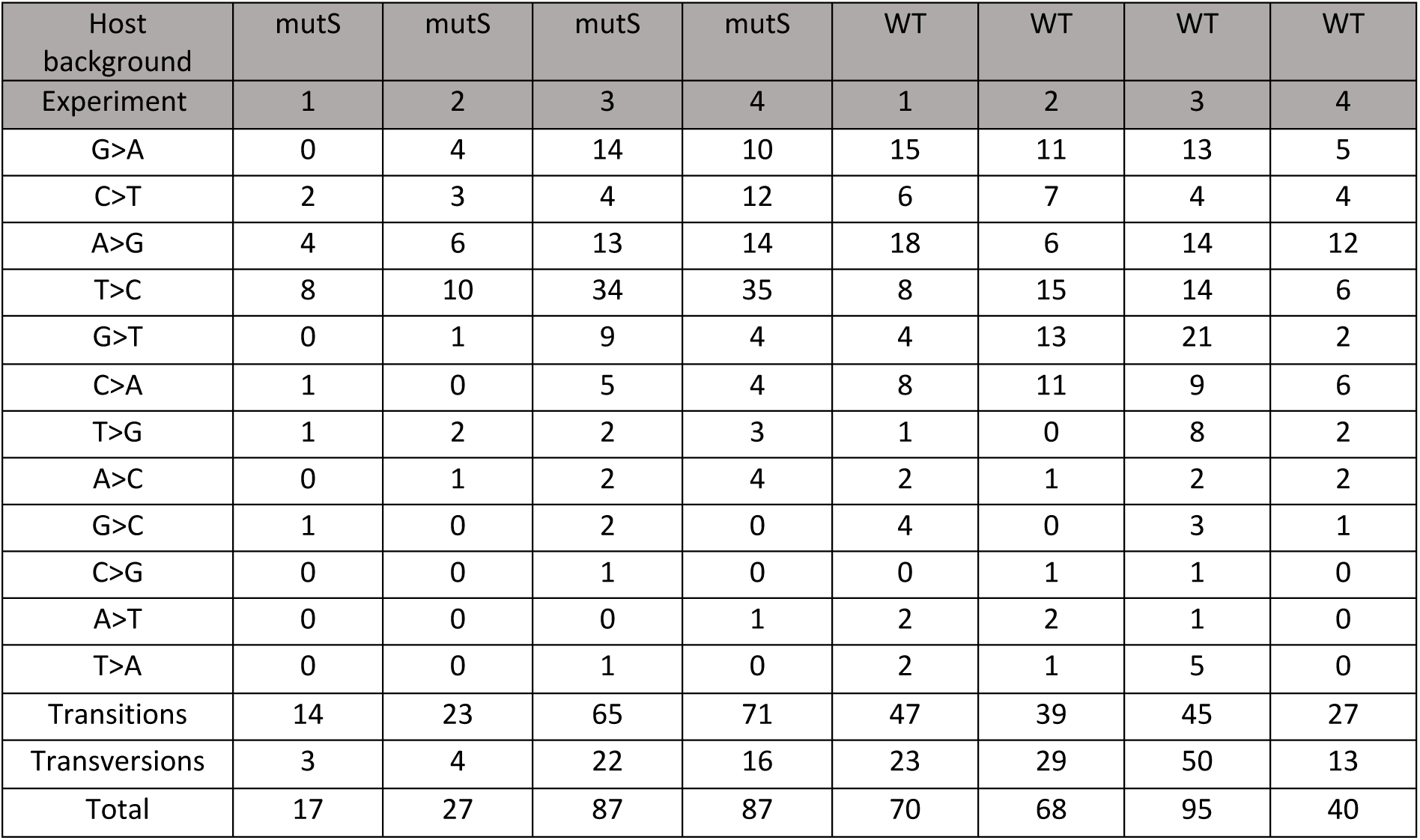
Spectrum of BPS leading to CII inactivation, as determined by *cII* Sanger sequencing. (full data are provided in Table S3)

| Host background | *mutS* | *mutS* | *mutS* | *mutS* | WT | WT | WT | WT |
| --- | --- | --- | --- | --- | --- | --- | --- | --- |
| Experiment | 1 | 2 | 3 | 4 | 1 | 2 | 3 | 4 |
| G>A | 0 | 4 | 14 | 10 | 15 | 11 | 13 | 5 |
| C>T | 2 | 3 | 4 | 12 | 6 | 7 | 4 | 4 |
| A>G | 4 | 6 | 13 | 14 | 18 | 6 | 14 | 12 |
| T>C | 8 | 10 | 34 | 35 | 8 | 15 | 14 | 6 |
| G>T | 0 | 1 | 9 | 4 | 4 | 13 | 21 | 2 |
| C>A | 1 | 0 | 5 | 4 | 8 | 11 | 9 | 6 |
| T>G | 1 | 2 | 2 | 3 | 1 | 0 | 8 | 2 |
| A>C | 0 | 1 | 2 | 4 | 2 | 1 | 2 | 2 |
| G>C | 1 | 0 | 2 | 0 | 4 | 0 | 3 | 1 |
| C>G | 0 | 0 | 1 | 0 | 0 | 1 | 1 | 0 |
| A>T | 0 | 0 | 0 | 1 | 2 | 2 | 1 | 0 |
| T>A | 0 | 0 | 1 | 0 | 2 | 1 | 5 | 0 |
| Transitions | 14 | 23 | 65 | 71 | 47 | 39 | 45 | 27 |
| Transversions | 3 | 4 | 22 | 16 | 23 | 29 | 50 | 13 |
| Total | 17 | 27 | 87 | 87 | 70 | 68 | 95 | 40 |

The modest increase in lambda’s mutation rate following MMR inactivation (Tables 1 & 2), along with the mutation spectrum distinct from that of MMR deficient *E. coli*, may reflect fundamental differences in the types of replication errors generated on the lambda genome versus the *E. coli* chromosome. To investigate this, we analyzed the mutation spectrum of lambda propagated in an MMR-deficient *E. coli* strain. Transitions accounted for only 79% of BPS in lambda compared to 97% in MMR deficient *E. coli* (Figure 4 & S6, Table 3, S3 & 4), indicating a substantial enrichment of transversions, mutations arising from errors that are much less efficiently repaired by MMR than those leading to transitions.

To further assess MMR activity on the lambda genome, we estimated the fold increase in mutation rates for transitions and transversions between MMR proficient and MMR deficient backgrounds. In *E. coli,* MMR deletion increases the transition rate by 240-fold and the transversion rate by only seven-fold (Lee et al. 2012). In contrast, on lambda, both categories showed an approximately six-fold increase in the absence of MMR, suggesting that MMR corrects transitions and transversions on the lambda genome with similarly moderate efficiency.

Together, these findings indicate an elevated frequency of transversion-type replication errors in lambda and suggest that MMR correct transitions less efficiently in lambda than in *E. coli*.

## Discussion

Mutation rate is a key parameter shaping the evolutionary potential of all organisms, as it determines how quickly new genetic variants arise and provides the raw material for natural selection. In bacteriophages, it critically influences phage-host dynamics by controlling how rapidly phages can adapt to bacterial defenses, evade resistance, and maintain long-term infectivity. A rate that is too low constrains adaptability, whereas a rate that is too high can impose a mutational load that reduces fitness. Precise knowledge of phage mutation rates is therefore essential not only for understanding page evolution but also for guiding their application in therapeutic contexts, i.e., phage therapy, the use of phages to treat bacterial infections. Phage therapy is challenged by the existence of numerous phage-resistant strains and variants that can be relatively easily selected within susceptible bacterial populations upon phage exposure. The ability of phages to adapt by evolving to counter bacterial resistance, is exploited in clinical settings by evolving phages on the targeted pathogen prior to their use in treatment, a procedure known as phage training. It increases the efficiency of treatment, by both improving the phage killing ability of the pathogen and preventing emergence of bacterial resistance in patients (Rohde et al. 2018; Pirnay et al. 2024). Phage mutation rate is a key factor in the dynamics of phage adaptation (Sanjuán and Domingo-Calap 2016), and therefore knowledge of this rate is crucial to predict phage adaptive capacity.

Yet very few studies determined the mutation rate of phages, and, to date, the only estimates of double-stranded DNA phage mutation rates were obtained on T4 (Santos and Drake 1994), T2 and lambda (reviewed in Drake 1991; Sanjuán et al. 2010), using FA on a single gene. In particular, the mutation rate of lambda was based on a single study (Dove 1968), in which Dove used a FA to estimate that inactivating mutations in the *cI* gene occur at a rate of 2 x10^-5^ per replication during the lytic cycle. However, the lack of raw data, experimental protocols, and calculation details limits the reliability of this estimate. Drake later analyzed two independently derived collections of *cI* inactivation mutants (Lieb 1981; Skopek and Hutchinson 1982) to estimate the fraction of mutations detectable by Dove’s assay and calculate what remained until now the only available per-base mutation rate for lambda, 7.7 x10^-8^. Because the detectable fraction varied substantially between the two collections, Drake estimated that lambda per-base mutation rate should be comprised between 2.7 ×10⁻^8^ and 2.2 ×10⁻^7^ (Drake 1991). Here we show that actual lambda mutation rate is 15-fold lower than its mean estimation, and at least five-fold lower than Drake’s lowest estimate. This difference might largely rely on the underestimation by Drake of the *cI* target size, as he hypothesized that among BPSs only those leading to stop codon were inactivating mutations. In our work, the sequencing of 854 cII mutants enabled to obtain a better estimation of the gene target size, as suggested by the similar mutation rate estimate obtained through FA and WGS.

Lambda phage itself has no therapeutic potential due to its temperate nature (its ability to stay in a prophage state in infected cells), but the results obtained and the methods developed with this model phage will be useful to establish the mutation rate of the strictly lytic (or virulent) phages used in therapy. Traditionally, phage mutation rates have been measured using FAs, which rely on detailed knowledge of phage biology and gene functions. However, such assays are not applicable to most therapeutic phages, for which this level of information is lacking. Here we show that DS is an easy to implement method suitable for obtaining a good approximation of the mutation rate of double-stranded DNA phages.

Although lower than previously estimated, the per-base mutation rate of lambda remains substantially higher, by approximately 20-fold, than that of its host, *E. coli,* despite both relying on the same replicative DNA polymerase. With the aim of explaining this difference, we investigated the mechanisms controlling lambda mutagenesis. While inactivation of many *E. coli* DNA repair systems has little or only a modest impact on mutation rate under non-stressful conditions, three pathways have been shown to play a major role in mutation control (Foster et al. 2015; Niccum et al. 2018). Inactivation of two of these pathways independently results in an approximately 150-fold increase in *E. coli*’s mutation rate (Foster et al. 2015; Niccum et al. 2018). We focused on one of these, the conserved MMR system, which corrects replication errors, and found that inactivation of host MMR only modestly increases lambda’s mutation rate, suggesting that MMR is significantly less effective on the lambda genome than on *E. coli* genome.

To investigate the basis of MMR’s low effectiveness on the lambda genome, we assessed both the ability of MMR to detect and repair replication errors in lambda DNA and the nature of replication errors generated during lambda infection. MMR is known to efficiently correct transition mutations but is markedly less effective at repairing transversions (Lee et al. 2012; Long et al. 2018). Our results suggest that replication error detection by MMR is not markedly compromised on the lambda genome. Moreover, we found no evidence that undermethylation of GATC sequences in lambda interferes with MMR activity, contradicting earlier hypothesis (Pukkila et al. 1983; Caillet-Fauquet et al. 1984; Caillet-Fauquet and Maenhaut-Michel 1988; Pereira-Gómez and Sanjuán 2015). Notably, comparison of the mutation spectra between lambda replicating in MMR deficient *E. coli* with those of the MMR-deficient *E. coli* revealed a significant enrichment in transversions in lambda. This finding is consistent with earlier observations by Wagner & Nohmi (Wagner and Nohmi 2000) and suggests that the distinctive replication error profile of lambda, marked by a higher frequency of transversion-causing errors, may underline the reduced MMR effect on lambda mutagenesis.

A possible explanation for the altered error spectrum observed in lambda is disruption in dNTP pool balance during infection. Imbalances in dNTP levels are known to increase mutation rates, likely by increasing the rate of replication errors or biasing the error spectrum (Kunz 1988; Bebenek et al. 1992). Consistent with this, gene expression profiling during lambda infection has revealed upregulation of the *E. coli* ribonucleotide reductase genes *nrdA*, *nrdB*, and *nrdD* (Osterhout et al. 2007; Liu et al. 2013). Overexpression of these genes has been associated with mutator phenotypes (Wheeler et al. 2005; Gon et al. 2006, 2011). dNTP pool imbalances have been reported during phage infection in *P. aeruginosa* (Chevallereau et al. 2016), suggesting that such metabolic shifts may be a general feature of phage replication.

In addition to a skewed replication error pattern, the limited effect of MMR on lambda mutagenesis may also stem from reduced activity of MutTMY system, another major *E. coli* repair system. Inactivation of this system leads to a mutation spectrum heavily dominated by specific transversions (∼99%) (Foster et al. 2015). Although this distinct signature is not present in our data, it was observed in an earlier study by Wagner & Nohmi (Wagner and Nohmi 2000), using the same reporter gene. The cause of this discrepancy remains unclear. Future studies should further investigate the mechanisms underlying the limited MMR correction of transitions in lambda, as well as the factors contributing to its atypical replication error pattern.

In addition to quantifying lambda’s mutation rate, our study reveals that inactivation of the bacterial MMR system can significantly modulate this rate, though to a markedly lesser extent than in *E. coli*, providing critical insight into the limits and potential of manipulating host repair pathways to accelerate phage evolution.

## Materials and Methods

### Strains and growth medium

For all experiments, we used the lambda ind1 *cI*_857_ (lambda PaPa *cI*_857 E118K_; see Sussman and Jacob 1962; Hendrix and Duda 1992) and its derivatives and wild-type *E. coli* MG1655 strain and its derivatives listed in Supplementary Table S8. Constructs of strains, phage and plasmids were verified through PCR, sequencing, and/or phenotypic assays, such as expression of fluorescent proteins or testing for the ability to generate mutations conferring resistance to rifampicin. Unless specified, all bacterial cultures were performed in LB broth (Bertani 1951) and phage infections done in LB + 10 mM MgSO_4_ + 0.2% maltose.

### Mutation accumulation experiments on lambda

30 parallel lines were obtained from a same phage stock (ancestral phage) and propagated for 150 MA cycles. Each cycle consisted in picking a lysis plaque, streaking it on a bacterial lawn and leaving it grow for 8 hours (day) or 16 hours (overnight). The top medium used was LB + Top Agarose (LB + 0.22% Agarose) supplemented with 10 mM MgSO_4_, 0.2% maltose and the bottom medium was LB Agar supplemented with either 25 µg/mL chloramphenicol or 50 µg/ml kanamycin depending on the bacterial strain. Every 9 cycles, one lysis plaque was purified by resuspension in LB and treatment with 10% chloroform and then stored at 4°C, then the next cycle started with this purification. This step prevented the appearance of phage-resistant and double-antibiotic resistant bacteria. On an average of 9 cycles, 12 lysis plaques were randomly chosen, resuspended and crushed in LB. The number of PFU per lysis plaque was determined by phage titration over the course of the MA experiment (4 duplicates). At cycle 150, one lysis plaque per line was resuspended and treated with chloroform. In order to prepare lysates for DNA extraction, these phages were amplified on a culture of *E. coli* WT at an OD_600_ of 0.08. The lysates were centrifugated 20 min at 4500g, filtered with a 0.2 μm filter, treated with Turbo DNase™ (Invitrogen™, US) and RNase I to remove the host contaminant DNA and RNA and precipitated with polyethylene glycol 10% and NaCl 0.5 M for 6 hours. The lysates were centrifuged and the pellets were resuspended in SM buffer, treated with proteinase K, and the phage DNA was purified using a standard phenol:chloroform procedure. DNA purity and concentration were checked with the NanoDrop™ 2000 spectrophotometer (Thermo Scientific™, US) and the Qubit4™ fluorometer (Invitrogen™, US), respectively. DNA integrity was checked by 1% agarose gel electrophoresis.

### Whole genome sequencing and analysis

Libraries were prepared by Eurofins for INVIEW resequencing. The reads were aligned on the reference genome NC_001416.1 from NCBI and variants were called using Breseq v0.38.1 (http://barricklab.org/brese^q^; see Deatherage and Barrick 2014). Variants that were identical to defective prophage Dlp12, Qin, Rac were excluded from the BPS analysis. Raw data are available at https://doi.org/10.57745/4RMO0D.

### Calculation of mutation rate from MA

The number of mutations was divided by the length of the genome (48,502), the number of lines (30), the number of cycles (150) and the number of DNA doublings per line. The last number was estimated from 74 measurements of the number of PFU per lysis plaque (see above), which is equal to 5.9 ± 0.3 x10^7^ PFUs per plaque.

### Amplification of lambda for Duplex Sequencing

Potential initial genetic heterogeneity within a lambda lysate was reduced by end-point dilutions. After the third dilution, 8 independent lines were started from 8 different plaques resuspended and incubated for 1 h at 4°C in 100 µL of SM buffer. After centrifugation to pellet bacteria, each supernatant was distributed on 4 plates (25 µl per plate), each sample mixed with 100 µl of a saturated culture of either WT or *mutS E. coli* and 3 ml of with Top Agar and poured onto 90 mm LB-Agar plates. The plates were incubated for 5h at 37°C until confluent lysis. Phages were recovered by pouring 6 mL of buffer (Tris-HCl pH 7.5 200 mM and 50 mM NaCl) on the plates, incubating overnight at 4°C and collecting together the buffer and the Top Agar from the 4 plates, that were vortexed and centrifuged for 15 min at 4500 g. The supernatants were filtered with a 0.2 μm filter and phage concentrations determined by plaque assay. 1 to 6 x10^11^ phage particles were obtained per line (mean of 4.4 x10^11^), which corresponds to 39 genome doublings assuming exponential replication. The phage DNA of the 8 lines was purified as described previously. In addition, plasmid pUC19 was purified, sequenced and analysed as a negative control. DNA purity and concentration were checked with the NanoDrop™ 2000 spectrophotometer (Thermo Scientific™, US) and the Qubit4™ fluorometer (Invitrogen™, US), respectively. DNA integrity was verified by 1% agarose gel electrophoresis.

### Sequencing of DS samples and analysis

Library preparation, sequencing and analysis were done as described previously (Risso-Ballester et al. 2016). Libraries were sequenced using a NextSeq™ 550 (Illumina®, US) sequencer yielding ∼100M paired-end reads. Raw reads passed through a Q20 filter in FastP v0.23.1. The filtered reads were analysed using the Duplex-seq-pipeline v2.1.4 (Kennedy et al. 2014) (https://github.com/Kennedy-Lab-UW/Duplex-Seq-Pipeline“ \o”https://github.com/Kennedy-Lab-UW/Duplex-Seq-Pipeline), the reference genome NC_001416.1 from NCBI and the per default run command except the family size which was reduced to 2. *E. coli* genome (NC_000913.3) was used in the pipeline as a contaminant genome to filter defective prophages reads that map on lambda genome. The mutations were extracted from the vcf files. Mutations present in all samples were filtered out (8 mutations in our ancestral lab strain), the others were used to determine the mutation spectrum. The mutation rate was determined by subtracting the number of mutations of each category in each sample by the number of mutations in the background pUC19. Then, the number of mutations of each type was divided by the number of sequenced bases, then by the estimated number of DNA doublings per sample (∼39). Raw reads are available at https://doi.org/10.57745/HROYNH.

### Determination of adenine methylation at GATC sites in lambda DNA

lambda phage was propagated on liquid cultures of *E. coli* strains expressing variable levels of Dam (depending on anhydrotetracycline concentration of 0, 5 or 50 µM), starting with a hundred phages (to make sure there was not a loss-of-function *cII* mutant in the inoculum) and 4 x10^5^ bacteria. Phage and bacteria were mixed in 10 µL of growth medium and incubated for 1 hour at 37°C. 1 ml of medium was then added, and the mixture incubated for 2 hours at 37°C, after which 9 ml of medium was added and the culture incubated with agitation until lysis occurred (monitored by a drop of optical density). The culture was centrifuged at 4,500 g for 10 minutes, the supernatant was filtered with a 0.2 µM size pore filter and stored at 4°C until use. Phage DNA was extracted as described previously, from 5 mL of lysate and sent for sequencing at the I2BC sequencing facility. DNA was sequenced with a GridION X5 (Oxford Nanopore Technologies, UK) and long-reads were analyzed with modkit v0.3.1 (https://github.com/nanoporetech/modki<u>^t^</u>) using per default parameters. The results of the comparative mode were used, using DNA from a dam knockout condition as reference (methylation level fixed at 0). Raw data are available at https://doi.org/10.57745/HIQF2N.

### Construction of lambda *cII*_G183A_ mutant

G at position 183 relative to the first base of *cII* gene was replaced by an A by Cas9 genome editing. To that aim, we constructed a plasmid, pMVRedcas9_cIIg183a, containing 3 different parts: i) the genes encoding the lambda Red recombination system proteins, ii) *cas9* and a 20 bp spacer targeting the WT *cII* sequence iii) the donor *cII* sequence containing the mutation (*cII*_G183A_). The 20 bp spacer was designed to be followed by a 5’-NGG-3’ sequence. The part i) was obtained by amplification of the plasmid pRed-cas9-Δpoxb300 (Zhao et al. 2016) with primers OMV310 containing the spacer at its 5’ end and OMV312. The fragment containing the *cII*_G183A_ sequence (iii) was obtained by site-directed mutagenesis. Three PCR reactions were performed using lambda WT DNA as template: the first 2 PCRs with primers OMV278-OMV279 and OMV280-OMV281 amplified the left and right parts of the target DNA, including the mutated region. The 3^rd^ PCR, carried out with primers OMV278 and OMV281, is an assembly PCR (double-joint PCR) of the 2 fragments as described in J. H. Yu et al., 2004 (Yu et al. 2004). The part i) was amplified with primers OMV304 and OMV314. All PCR fragments were gel purified using the Nucleospin® PCR and gel cleanup kit (Macherey-Nagel, DE) with plasmid PCRs first digested by DpnI to eliminate the template. The 3 parts were then assembled by Gibson assembly using the NEBuilder® HiFi DNA Assembly Master Mix (New England Biolab™, US), using overlapping primers OMV304 and OMV314 for part i), OMV311 and OMV312 for part ii) and OMV308 and OMV309 for part iii). The Gibson mixture was then electroporated into a *E. coli* DH10B strain. Kanamycin resistant clones were streaked and cultivated at 30°C and plasmid extracted with NucleoSpin Plasmid DNA extraction kit (Macherey Nagel). The sequence of the plasmid pMVRedcas9-cIIg183a was verified by WPS at Eurofins. The lysogenic strain MVEC276 *λ_cI857_ ΔrecA* used (constructed by transduction with a P1 from strain GSY5902), to prevent lambda excision due to a SOS response induced by Cas9 activity, was then electroporated with the resulting plasmid. Cas9 and lambda Red were induced as described in (Zhao et al. 2016) by cultivating the transformants in LB + 0.2% L-arabinose for 6 hours at 30°C, followed by plating onto LBA plates supplemented with 50 µg/mL kanamycin and 0.2% L-arabinose to select recombinants. The presence of the *cII* G183A mutation was verified by Sanger sequencing (TubeSeq Supreme from Eurofins Genomics, DE) on PCRs using primers cII_up and cII_dwn. The lambda prophage was then thermally induced and phage used to lysogenize strain MG1655. Clones selected for their thermosensitivity were checked for monolysogeny by PCR using primers JM50, JM51 and JM52 (Table S9), then 2 clones were stored (MV318 and MV319). Phage stocks G183A_L1 and G183A_L2 were produced by induction of strains MV318 and MV319 respectively.

### Construction of MVEC337 strain

Strain LZ613 (Trinh et al., 2020), carrying prophage λ*cI*_857_ *bor*::*KanR*, was transformed with plasmid pLZ1 (Trinh et al., 2020). Heat-shock induction of the prophage allowed recombination with pLZ1 during replication, generating λ*cI*857 *bor::cat-* 21tetO phages. These recombinants were used to lysogenize strain MG1655 6300 using chloramphenicol selection and the resulting lysogen served to produce a lysate of lambda *cI*857 *bor*::*cat* 21tetO phage. Strain MVEC337, used for microscopy, was obtained by lysogenizing MVEC333 (expressing YFP-MutL), with this phage, and subsequently transformation it with the plasmid pLZ3 (Trinh et al., 2020), which expresses TetR-mCherry

### Fluctuation assay on lambda *cII*

Two independent lambda phage stocks (WT_L1 and WT_L2) were prepared by induction of *E. coli* MD114 and MVEC235 (see Table S8), lysogenic for lambda phage. The fluctuation assay was performed in single burst with these stocks, as well as with phage stocks G183A_L1 and G183A_L2, using either WT or *mutS E. coli* cells, and the *cII* selection system (Jakubczak et al. 1996). At least 3 assays were performed per strain and per phage genotype, using at least 20 parallel infected cultures. 6 additional cultures per assay were performed, 3 for estimation of the initial number of phages (*N_0_*, before growth on the non-selective strain, estimated without selection), 3 for the estimate of the final number of phages (*N_1_*, after growth on the non-selective strain and without selection, using FtsH-cells at 37°C). To obtain infected cells, 10 µL of phage at an m.o.i. of 1 were added to 1 mL of *E. coli* culture at an OD600 of 0.25 grown at 37°C and incubated fir 10 min at 37°C. Cultures were then diluted in LB (50-fold for WT, 500-fold for *mutS*, and 5-fold less for G183A stocks) and distributed into 1.5 mL microtubes (100 µL per tube). After incubation at 37°C for 1h, the samples were mixed with 30 µL of a saturated selective culture (FtsH-deficient *E. coli)*, mixed with 1 ml of Top Agarose and poured on 60 mm diameter LBA plates. Lysis plaques were counted after 48H of incubation at 25°C.

### Determination of lambda mutation rate and mutation spectrum with fluctuation assay

The number of mutants per culture and the *P_0_* fraction were determined for each assay. The mutation rate per target (*cII*) per replication was estimated using the MSS-MLE method (Ma et al. 1992) with the webtool SALVADOR (https://websalvador.eeeeeric.com<u>^/^</u>, see Zheng 2002, 2021) and the null-class (*P_0_*) estimator (see Luria and Delbrück 1943). Plating efficiency of the selective cells was estimated as 0.58, but was not used as *N_1_* was determined using the selective strain at 37°c. To determine the mutation spectrum, *cII* genes from lysis plaques on selective FtsH-hosts were amplified and sequenced. In addition to mutants obtained in the fluctuation assay, other mutants were obtained by simply by distributing 10 µL of infected cultures (obtained as described above) in 96-well plates. After incubation for 1h at 37°C, the whole 10µL cultures were spotted on a layer of FtsH-deficient *E. coli* cells (in Top Agarose, see above) on 12 cm-side square LBA plates. One lysis plaque per independent culture was used for *cII* amplification by PCR using OneTaq® DNA polymerase (New England Biolab™, US) and manufacturer’s instructions with an initial denaturation step at 95°C of 5 min, using primers cII_up and cII_dwn (Table S9). Unpurified PCR product size and concentration was checked with 1% agarose gel electrophoresis and sequenced by PlateSeq Supreme Sanger sequencing (Eurofins Genomics, DE), using cII_up as primer. Mutations were identified by visualization of the traces mapped on *cII* reference sequence (NC_001416.1: 38360-38653) in Clone Manager v10.1 (https://scied.com/index.ht<u>^m^</u>). BPS and indels rate per base replicated were estimated as described by G. I. Lang & A. W. Murray (Lang and Murray 2008), by dividing the individual BPS and indels rate per locus by their respective target size of BPS (134.8 for WT, 143.6 for *mutS*) and indels (294 = length of *cII* sequence). The rate was corrected for the T>G BPS in position 185, linked to the 6G hotspot (7.4% for WT, 1.3% for *mutS*, BPS not observed with the G183A phages). Raw data are available at https://doi.org/10.57745/TH0GZF.

### Microfluidic experiments

The mother machine device is described in Robert et al. (Robert et al. 2018). It consists of a main trench 15mm long, 50µm wide and ∼30µm high and ∼1000 perpendicular microchannels which are ∼25µm long and ∼1µm wide and high. To produce PDMS chips from the molds we followed the protocol indicated in L. Robert et al. (Robert et al. 2018, 2019). For delivering and controlling the flow of growth medium into the chip, we used a PHD Ultra™ syringe pump (Harvard Apparatus™, US) set at 2ml/h, 50 ml Monoject™ syringes with LS23 needles (Phymep, FR), Tygon S54-HL flexible tubing (Phymep, FR) and SC23/8 steel couplers (Phymep, FR). All experiments were performed with LB as a growth medium. To induce *yfp-*mutL expression, the medium was supplemented with 1mM IPTG.

### Microscopy

Fluorescence imaging was performed as in L. Robert et al. (Robert et al. 2018) with a Delta Vision Elite inverted microscope (Image Solutions, UK) equipped with the Ultimate Focus system for automatic focalization (Image Solutions, UK), a 100x oil immersion objective (N.A. 1.4: Olympus), a temperature-controlled chamber (Image Solutions, UK), and the DV Elite sCMOS camera (Image Solutions, UK). Fluorescence illumination was provided by the DV Light Solid-State Illuminator 7 Colours (Image Solutions, UK) at 575 nm (mCherry excitation) and 513 nm (YFP excitation). The temperature controller was set to 36.5°C. The waiting time before acquisition was 3 hours to allow the cells to adapt to the environment and reach stable exponential growth. During acquisitions, the temperature was either maintained at 36.5°C, for the control condition, i.e., *E. coli* cells carrying phage lambda *cI_857_* in the lysogenic state, or increased to 39.5°C to induce the lytic cycle, with data acquisition starting 15 minutes after the temperature shift. On average, 10 fields of view were selected per experiment, with an average of ∼15 microchannels per field of view. Images were acquired every Δt = 2 minutes for ∼3h. For mCherry illumination, we used an exposure time of 0.2 second and a maximum LED intensity of 3%, and for YFP illumination, an exposure time of 2 seconds and a maximum LED intensity of 7%. As in Robert et al. (Robert, Science, 2018), for YFP images, we used the optical axis integration imaging mode, that allows collecting fluorescent light by integrating one image through a z-axis movement of 1 µm around the cell focal plane. All experiments were repeated at least twice.

### Image analysis

Images were analysed using BACMMAN v3.8.4 (https://github.com/jeanollion/bacmma^n^), a custom ImageJ plugin (https://github.com/imagej/ImageJ) (Ollion et al. 2019; Robert et al. 2019). Segmentation and tracking of the microchannels over time were performed using mCherry images and the *MicrochannelFluo2D* and *MicrochannelTracker* modules. Bacteria segmentation and tracking from mCherry images were performed using the *BacteriaFluo* and *BacteriaClosedMicrochannelTrackerLocalCorrections* modules. YFP-MutL foci were segmented and tracked over time from YFP images using the *SpotSegmenterRS3D*, and *NestedSpotTracker* modules. BACMMAN automatically measures characteristics of bacteria (e.g., growth rate, cell size and length) and foci (e.g., lifetime and time of onset).

### Data analysis

Data analysis was performed using custom codes developed in R language.

For DS data, we estimated the cumulative mutation frequency after 39 DNA doublings at the genome wide level (respectively in sliding windows of 1kb) assuming that mutations arise according to a homogeneous Poisson process: the number of mutations was divided by the number of sequenced bases. 95% confidence intervals for each window were then computed based on the number of sequenced bases to assess whether the local estimates were compatible with the genome level one. Finally, the distance between consecutive mutations was compared to the exponential distribution (theoretical if the mutation rate is constant) using a chi-square goodness of fit test after binning distances in bins of length 0.5kb.

For YFP-MutL visualization data the mean rate of YFP-MutL foci was calculated as the mean number of newly appeared foci per “cell track”, where a “cell track” is the time between birth and division for lysogenic *E. coli* at 35°C and non-lysogenic *E. coli* (experiments: WT 𝞴 hom 35°C, WT 𝞴 35°C, WT 37°C in Table S10), and from birth to lysis for lysing cells (experiments: WT 𝞴 39°C, WT 𝞴 mutS 39°C, WT 𝞴 hom 39°C in Table S10). Foci present at frame 0 were excluded from the analysis. For the experiments at 36.5°C and 37°C, the analysis was restricted to mother cells with complete tracks, i.e., mother cells tracked from birth to division. Only YFP-MutL foci from these non-truncated tracks were included in the rate calculations. In the 39°C experiments the analysis was limited to the lysing cells, regardless of whether they were mother or daughter cells to increase statistical power, except for the experiment WT lambda 39°C shown in Figure 2d, e, f, for which only mother cells were considered. The experiment WT 37°C (from Enrico Bena et al. 2024) was acquired over an extended period of time (46 hours, 11997 mother cell tracks). To calculate the rate (Table S10) and the number of newly appeared YFP-MutL foci per cell (Figure 2E), we applied a bootstrapping approach. Specifically: i) we performed 300 resampling, each involving 600 cell tracks randomly sampled with replacement from the original dataset, ii) for each resampled set, we computed the mean YFP-MutL foci rate as previously described, and iii) we then calculated the overall mean and +/2*SEM from the distribution of these 300 mean rates. For the data shown in Figure 2d, we applied a bootstrapping approach as follows: i) 300 resampling were performed, each involving the random selection (with replacement) of 120 non-truncated mother cell lineages, each truncated to 66 minutes, from the original dataset. This truncation and sample size were chosen to allow comparison with the WT λ 36.5 °C and *mutS* λ 39 °C experiments. ii) For each of 300 datasets, we calculated the mean number of newly appeared YFP-MutL foci per frame. Iii) For each time point, we then computed the average of these means across the 300 resamples, along with +/2*SEM. Note that in Table S10, the average rate for WT λ 35 °C is approximately twice that of WT λ hom 35 °C. This difference is expected, as the two experiments were performed with different time resolutions, 1 minute and 2 minutes, respectively (Enrico Bena et al. 2024).

### Quantitative PCR (qPCR)

Overnight cultures of lysogenic bacterial cultures were diluted 200-fold in 10 ml of LB and incubated with agitation at 35°C. At a OD600 of 0.4, cultures were shifted to 42°C for 10 minutes, and then incubated at 37°C. Just before and after the thermal shift as well as after 15, 30 and 45 minutes at 37°C, 1 ml aliquots of culture were centrifuged for 2 minutes at 8,000 rpm. Supernatant was carefully removed and the pellet frozen at-20°C. Pellets were resuspended in 100 μL of resuspension solution (50 mM Tris-HCl (pH 8.0), 50mM glucose, 50mM NaCl, 2 mg/ml lysozyme) and incubated for 10 min at room temperature. 100 μL of lysis buffer (50 mM Tris-HCl (pH 8.0), 1% NP-40, 100 μg/ml RNAse, 10 mM EDTA, 50 mM NaCl) was added and incubated on ice for 30 min. Samples were then incubated with 0,5 % SDS and 50 μg/ml proteinase K for 1 hour at 65°C. DNA was purified using NucleoSpin Tissue XS kit (Macherey Nagel) following manufacturer’s instructions. *E. coli* and lambda genome copy numbers were assessed by qPCR on *gyrB* and *Q* genes respectively, using primers in Table S9. qPCR was performed in a total volume of 15 µL in MicroAmp Fast Optical 96-well plates sealed with MicroAmp Optical Adhesive Film using the Takyon ROX SYBR Mastermix blue dTTP kit. Amplifications were run in duplicate on a StepOnePlus real-time PCR system with the following cycling conditions: 95°C for 5 min, (95°C for 15 s, 58°C for 45 s, 72°C for 30 s) for 45 cycles, 72°C for 5 min, 95°C for 15 s, 60°C for 15 s, 95°C for 15 s. The relative lambda genome copy number was estimated by the ΔCT method: *Q* to *gyrB* copy number was set to one in the prophage state (before induction), which enabled to determine the *Q* to *gyrB* ratio during the lytic cycle. Mean *E. coli* genome copy number per cell was hypothesized to be 3, based on the CCSim tool (https://sils.fnwi.uva.nl/bcb/cellcycle/), which enabled to estimate the mean lambda copy number per cell at the different time points.

## Acknowledgments

We thank Dr. José M. Cuevas and the Genomics Facility of the Universitat de València (Spain) for DupSeq sequencing, Delphine Naquin and Yan Jaszczyszyn from the Next Generation Sequencing Core Facility of the I2BC Institute (Gif-sur-Yvette, France) for nanopore sequencing, and Agnès Thierry (Institut Pasteur) for Illumina sequencing. We are grateful to Lanying Zeng for providing λ*cI*_857_ *bor*::*KanR* and associated plasmids, and to Jean Ollion (SABILab) for image analysis tools. We are grateful to the INRAE MIGALE bioinformatics facility (MIGALE, INRAE, 2020. Migale bioinformatics Facility, doi:10.15454/1.5572390655343293E12) for providing help, computing, and storage resources. We acknowledge the sequencing and bioinformatics expertise of the I2BC High-throughput sequencing facility, supported by France Génomique (funded by the French National Program “Investissement d’Avenir” ANR-10-INBS-09).

## References

Bebenek K, Roberts JD, Kunkel TA (1992) The effects of dNTP pool imbalances on frameshift fidelity during DNA replication. J Biol Chem 267:3589–3596. 10.1016/s0021-9258(19)50565-8

Bednarz M, Halliday JA, Herman C, Golding I (2014) Revisiting bistability in the lysis/lysogeny circuit of bacteriophage lambda. PLoS One 9:. 10.1371/journal.pone.0100876

Bertani G (1951) Studies on lysogenesis. I. The mode of phage liberation by lysogenic Escherichia coli. J Bacteriol 62:293–300

Caillet-Fauquet P, Maenhaut-Michel G (1988) Nature of the SOS mutator activity: Genetic characterization of untargeted mutagenesis in Escherichia coli. MGG Mol Gen Genet 213:491–498. 10.1007/BF00339621

Caillet-Fauquet P, Maenhaut-Michel G, Radman M (1984) SOS mutator effect in E. coli mutants deficient in mismatch correction. EMBO J 3:707–712. 10.1002/j.1460-2075.1984.tb01873.x

Campbell JL, Kleckner N (1988) The rate of Dam-mediated DNA adenine methylation in Escherichia coli. Gene 74:189–190. 10.1016/0378-1119(88)90283-1

Chevallereau A, Blasdel BG, De Smet J, et al (2016) Next-Generation “-omics” Approaches Reveal a Massive Alteration of Host RNA Metabolism during Bacteriophage Infection of Pseudomonas aeruginosa. PLoS Genet 12:1–20. 10.1371/journal.pgen.1006134

De Paepe M, Hutinet G, Son O, et al (2014) Temperate Phages Acquire DNA from Defective Prophages by Relaxed Homologous Recombination: The Role of Rad52-Like Recombinases. PLoS Genet 10:. 10.1371/journal.pgen.1004181

Deatherage DE, Barrick JE (2014) Identification of Mutations in Laboratory-Evolved Microbes from Next-Generation Sequencing Data Using breseq. In: Sun L, Shou W (eds) Engineering and Analyzing Multicellular Systems: Methods and Protocols, Methods in Molecular Biology. Springer Science+Business Media, New York, pp 165–188

Dillon MM, Sung W, Lynch M, Cooper VS (2018) Periodic variation of mutation rates in bacterial genomes associated with replication timing. MBio 9:1–15. 10.1128/mBio.01371-18

Dillon MM, Sung W, Lynch M, Cooper VS (2015) The rate and molecular spectrum of spontaneous mutations in the GC-rich multichromosome genome of Burkholderia cenocepacia. Genetics 200:935–946. 10.1534/genetics.115.176834

Dillon MM, Sung W, Sebra R, et al (2017) Genome-wide biases in the rate and molecular spectrum of spontaneous mutations in vibrio cholerae and vibrio fischeri. Mol Biol Evol 34:93–109. 10.1093/molbev/msw224

Dohet C, Wagner R, Radman M (1986) Methyl-directed repair of frameshift mutations in heteroduplex DNA. Proc Natl Acad Sci U S A 83:3395–3397. 10.1073/pnas.83.10.3395

Dove WF (1968) The genetics of the lambdoid phages. Annu Rev Genet 305–340

Drake JW (1991) A constant rate of spontaneous mutation in DNA-based microbes. Proc Natl Acad Sci U S A 88:7160–7164. 10.1073/pnas.88.16.7160

Drake JW, Charlesworth B, Charlesworth D, Crow JF (1998) Rates of spontaneous mutation. Genetics 148:1667–1686. 10.1093/genetics/148.4.1667

Dreiseikelmann B, Eichenlaub R, Wackernagel W (1979) The effect of differential methylation by Escherichia coli of plasmid DNA and phage T7 and lambda DNA on the cleavage by restriction endonuclease MboI from Moraxella bovis BRIGITTE. Biochem Biophys Res Commun 562:418–428

Duffy S, Shackelton LA, Holmes EC (2008) Rates of evolutionary change in viruses: Patterns and determinants. Nat Rev Genet 9:267–276. 10.1038/nrg2323

Enrico Bena C, Ollion J, De Paepe M, et al (2024) Real-time monitoring of replication errors’ fate reveals the origin and dynamics of spontaneous mutations. Nat Commun 15:. 10.1038/s41467-024-46950-0

Foster PL (2006) Methods for Determining Spontaneous Mutation Rates. Methods Enzym 195–213

Foster PL, Hanson AJ, Lee H, et al (2013) On the mutational topology of the bacterial genome. G3 Genes, Genomes, Genet 3:399–407. 10.1534/g3.112.005355

Foster PL, Lee H, Popodi E, et al (2015) Determinants of spontaneous mutation in the bacterium Escherichia coli as revealed by whole-genome sequencing. Proc Natl Acad Sci U S A 112:E5990–E5999. 10.1073/pnas.1512136112

Glickman BW, Radman M (1980) Escherichia coli mutator mutants deficient in methylation-instructed DNA mismatch correction. Proc Natl Acad Sci U S A 77:1063–1067. 10.1073/pnas.77.2.1063

Gon S, Camara JE, Klungsøyr HK, et al (2006) A novel regulatory mechanism couples deoxyribonucleotide synthesis and DNA replication in Escherichia coli. EMBO J 25:1137– 1147. 10.1038/sj.emboj.7600990

Gon S, Napolitano R, Rocha W, et al (2011) Increase in dNTP pool size during the DNA damage response plays a key role in spontaneous and induced-mutagenesis in Escherichia coli. Proc Natl Acad Sci U S A 108:19311–19316. 10.1073/pnas.1113664108

Hattman S (1972) Plasmid-Controlled Variation in the Content of Methylated Bases in Bacteriophage Lambda Deoxyribonucleic Acid. J Virol 10:356–361. 10.1128/jvi.10.3.356-361.1972

Hénaut A, Rouxel T, Gleizes A, et al (1996) Uneven distribution of GATC motifs in the Escherichia coli chromosome, its plasmids and its phages. J Mol Biol 257:574–585. 10.1006/jmbi.1996.0186

Hendrix RW, Duda RL (1992) Bacteriophage lambda PaPa: Not the Mother of All lambda Phages. Science (80-) 258:1145–1148

Jakubczak JL, Merlino G, French JE, et al (1996) Analysis of genetic instability during mammary tumor progression using a novel selection-based assay for in vivo mutations in a bacteriophage λ transgene target. Proc Natl Acad Sci U S A 93:9073–9078. 10.1073/pnas.93.17.9073

Kennedy SR, Schmitt MW, Fox EJ, et al (2014) Detecting ultralow-frequency mutations by Duplex Sequencing. Nat Protoc 9:2586–2606. 10.1038/nprot.2014.170

Kunz BA (1988) Mutagenesis and deoxyribonucleotide pool imbalance. Mutat Res - Fundam Mol Mech Mutagen 200:133–147. 10.1016/0027-5107(88)90076-0

Lang GI, Murray AW (2008) Estimating the per-base-pair mutation rate in the yeast Saccharomyces cerevisiae. Genetics 178:67–82. 10.1534/genetics.107.071506

Lee H, Popodi E, Tang H, Foster PL (2012) Rate and molecular spectrum of spontaneous mutations in the bacterium Escherichia coli as determined by whole-genome sequencing. Proc Natl Acad Sci U S A 109:. 10.1073/pnas.1210309109

Leong PM, Hsia HC, Miller JH (1986) Analysis of spontaneous base substitutions generated in mismatch-repair-deficient strains of Escherichia coli. J Bacteriol 168:412–416. 10.1128/jb.168.1.412-416.1986

Lieb M (1981) A fine structure map of spontaneous and induced mutations in the lambda repressor gene, including insertions of IS elements. MGG Mol Gen Genet 184:364–371. 10.1007/BF00352506

Liu X, Jiang H, Gu Z, Roberts JW (2013) High-resolution view of bacteriophage lambda gene expression by ribosome profiling. Proc Natl Acad Sci U S A 110:11928–11933. 10.1073/pnas.1309739110

Long H, Miller SF, Williams E, Lynch M (2018) Specificity of the DNA mismatch repair system (MMR) and mutagenesis bias in bacteria. Mol Biol Evol 35:2414–2421. 10.1093/molbev/msy134

Long H, Sung W, Miller SF, et al (2014) Mutation rate, spectrum, topology, and context-dependency in the DNA mismatch repair-deficient Pseudomonas fluorescens ATCC948. Genome Biol Evol 7:262–271. 10.1093/gbe/evu284

Luria SE, Delbrück M (1943) Mutations of Bacteria From Virus Sensitivity To Virus Resistance. Genetics 28:491–511. 10.1093/genetics/28.6.491

Lynch M (2010) Evolution of the mutation rate. Trends Genet 26:345–352. 10.1016/j.tig.2010.05.003

Lynch M, Ackerman MS, Gout JF, et al (2016) Genetic drift, selection and the evolution of the mutation rate. Nat Rev Genet 17:704–714. 10.1038/nrg.2016.104

Ma WT, Sandri G vH., Sarkar S (1992) Analysis of the Luria-Delbrück distribution using discrete convolution power. J Appl Prob 29:255–267

Marinus MG, Morris NR (1975) Pleiotropic effects of a DNA adenine methylation mutation (dam-3) in Escherichia coli K12. Mutat Res 28:15–26. 10.1016/0027-5107(75)90309-7

Niccum BA, Lee H, MohammedIsmail W, et al (2018) The spectrum of replication errors in the absence of error correction assayed across the whole genome of escherichia coli. Genetics 209:1043–1054. 10.1534/genetics.117.300515

Ollion J, Elez M, Robert L (2019) High-throughput detection and tracking of cells and intracellular spots in mother machine experiments. Nat Protoc 14:3144–3161. 10.1038/s41596-019-0216-9

Osterhout RE, Figueroa IA, Keasling JD, Arkin AP (2007) Global analysis of host response to induction of a latent bacteriophage. BMC Microbiol 7:1–12. 10.1186/1471-2180-7-82

Pereira-Gómez M, Sanjuán R (2015) Effect of mismatch repair on the mutation rate of bacteriophage /X174. Virus Evol 1:1–9. 10.1093/ve/vev010

Pirnay JP, Djebara S, Steurs G, et al (2024) Personalized bacteriophage therapy outcomes for 100 consecutive cases: a multicentre, multinational, retrospective observational study. Nat Microbiol 9:1434–1453. 10.1038/s41564-024-01705-x

Pukkila PJ, Peterson J, Herman G, et al (1983) Effects of high levels of DNA adenine methylation on methyl-directed mismatch repair in Escherichia coli. Genetics 104:571–582. 10.1093/genetics/104.4.571

Risso-Ballester J, Cuevas JM, Sanjuán R (2016) Genome-Wide Estimation of the Spontaneous Mutation Rate of Human Adenovirus 5 by High-Fidelity Deep Sequencing. PLoS Pathog 12:1–16. 10.1371/journal.ppat.1006013

Robert L, Ollion J, Elez M (2019) Real-time visualization of mutations and their fitness effects in single bacteria. Nat Protoc 14:3126–3143. 10.1038/s41596-019-0215-x

Robert L, Ollion J, Robert J, et al (2018) Mutation dynamics and fitness effects followed in single cells. Science (80-) 359:1283–1286. 10.1126/science.aan0797

Rohde C, Resch G, Pirnay JP, et al (2018) Expert opinion on three phage therapy related topics: Bacterial phage resistance, phage training and prophages in bacterial production strains. Viruses 10:. 10.3390/v10040178

Sanjuán R, Domingo-Calap P (2016) Mechanisms of viral mutation. Cell Mol Life Sci 73:4433– 4448. 10.1007/s00018-016-2299-6

Sanjuán R, Nebot MR, Chirico N, et al (2010) Viral Mutation Rates. J Virol 84:9733–9748. 10.1128/jvi.00694-10

Santos ME, Drake JW (1994) Rates of Spontaneous Mutation in Bacteriophage T4 Are Independent. Genetics 138:553–564

Schaaper RM, Dunn RL (1987) Spectra of spontaneous mutations in Escherichia coli strains defective in mismatch correction: the nature of in vivo DNA replication errors. Proc Natl Acad Sci U S A 84:6220–6224. 10.1073/pnas.84.17.6220

Schmitt MW, Kennedy SR, Salk JJ, et al (2012) Detection of ultra-rare mutations by next-generation sequencing. Proc Natl Acad Sci U S A 109:14508–14513. 10.1073/pnas.1208715109

Sergueev K, Court D, Reaves L, Austin S (2002) E. coli cell-cycle regulation by bacteriophage lambda. J Mol Biol 324:297–307. 10.1016/S0022-2836(02)01037-9

Skopek TR, Hutchinson F (1982) DNA base sequence changes induced by bromouracil mutagenesis of lambda phage. J Mol Biol 159:19–33. 10.1016/0022-2836(82)90029-8

Sussman R, Jacob F (1962) [On a thermosensitive repression system in the Escherichia coli lambda bacteriophage]. C R Hebd Seances Acad Sci 254:1517–1519

Szyf M, Avraham-Haetzni K, Reifman A (1984) DNA methylation pattern is determined by the intracellular level of the methylase. Proc Natl Acad Sci U S A 81:3278–3282. 10.1073/pnas.81.11.3278

Taylor K, Wȩgrzyn G (1995) Replication of coliphage lambda DNA. FEMS Microbiol Rev 17:109–119. 10.1016/0168-6445(95)00077-1

Trinh JT, Shao Q, Guan J, Zeng L (2020) Emerging heterogeneous compartments by viruses in single bacterial cells. Nat Commun 11:1–11. 10.1038/s41467-020-17515-8

Wagner J, Nohmi T (2000) Escherichia coli DNA polymerase IV mutator activity: Genetic requirements and mutational specificity. J Bacteriol 182:4587–4595. 10.1128/JB.182.16.4587-4595.2000

Wheeler LJ, Rajagopal I, Mathews CK (2005) Stimulation of mutagenesis by proportional deoxyribonucleoside triphosphate accumulation in Escherichia coli. DNA Repair (Amst) 4:1450–1456. 10.1016/j.dnarep.2005.09.003

Wold MS, Mallory JB, Roberts JD, et al (1982) Initiation of bacteriophage λ DNA replication in vitro with purified λ replication proteins. Proc Natl Acad Sci U S A 79:6176–6180. 10.1073/pnas.79.20.6176

Yu JH, Hamari Z, Han KH, et al (2004) Double-joint PCR: A PCR-based molecular tool for gene manipulations in filamentous fungi. Fungal Genet Biol 41:973–981. 10.1016/j.fgb.2004.08.001

Zhao D, Yuan S, Xiong B, et al (2016) Development of a fast and easy method for Escherichia coli genome editing with CRISPR/Cas9. Microb Cell Fact 15:1–9. 10.1186/s12934-016-0605-5

Zheng Q (2015) Methods for comparing mutation rates using fluctuation assay data. Mutat Res - Fundam Mol Mech Mutagen 777:20–22. 10.1016/j.mrfmmm.2015.04.002

Zheng Q (2021) webSalvador: a Web Tool for the Luria-Delbrük Experiment. Microbiol Resour Announc 10:6–8. 10.1128/mra.00314-21

Zheng Q (2002) Statistical and algorithmic methods for fluctuation analysis with SALVADOR as an implementation. Math Biosci 176:237–252. 10.1016/S0025-5564(02)00087-1

Zou X, Koh GCC, Nanda AS, et al (2021) A systematic CRISPR screen defines mutational mechanisms underpinning signatures caused by replication errors and endogenous DNA damage. Nat Cancer 2:643–657. 10.1038/s43018-021-00200-0

Zylicz M, Ang D, Liberek K, Georgopoulos C (1989) Initiation of λ DNA replication with purified host-and bacteriophage-encoded proteins: The role of the dnaK, dnaJ and grpE heat shock proteins. EMBO J 8:1601–1608. 10.1002/j.1460-2075.1989.tb03544.x

