## Supplementary Information for "Atypical Replication Error Pattern and Limited Repair Efficiency Contribute to Elevated Mutation Rate in Phage lambda"

Affiliation:
[1]Université Paris-Saclay, INRAE, AgroParisTech, Micalis Institute, Jouy-en-Josas, France
[2]Université Paris-Saclay, INRAE, MaIAGE, Jouy-en-Josas, France
[3]Université Paris-Saclay, INRAE, BioinfOmics, MIGALE bioinformatics facility, Jouy-en- Josas, France.
[4]Virus Evolution Laboratory, Institute for Integrative Systems Biology (I2SysBio), Universitat de València-CSIC, Paterna, Spain

**SUPPLEMENTARY INFORMATION**

### Index

### Supplementary text

**1.1 Estimating the number of mutations per lambda lytic cycle in the absence of error repair.** The lambda DNA is replicated by *E. coli* DNA polymerase III (Taylor and Węgrzyn, 1995). To estimate the number of mutations per lytic cycle, under the assumption that none of the replication errors are corrected, we multiplied the total number of bases copied by the mutation rate observed in MMR deficient *E. coli*, as in these cells, all replication errors introduced by DNA polymerase III are converted into mutations (Lee et al., 2012). During a typical infection, a lambda phage injects its 48,502 bp genome into an *E. coli* cell and produces ~250 progeny. Because replication is semi-conservative, this results in the synthesis of 498 new DNA strands, totaling ~24 million bases copied per lytic cycle. Applying a mutation rate of $3.26 \pm 1.51 \times 10^{-8}$ mutations per base, we estimate that each lytic cycle produces approximately 0.95 mutations, assuming no correction of replication errors.

**1.2 Estimating the number of mutations per lambda lytic cycle from YFP-MutL foci.** To estimate the number of mutations per infection cycle based on the observed number of YFP-MutL foci, which mark replication errors, we accounted for the segregation of the mismatch-containing DNA molecule and assumed no repair occurs, as detailed in (Robert et al. 2018). Briefly, each round of replication produces two double-stranded DNA molecules, four strands in total, and only one strand carries the replication error. Thus, a single mismatch containing DNA (marked by a YFP-MutL focus) results in one mutant genome and 3 wild-type genomes. Given an average of 11.1 ± 0.9 YFP-MutL foci per infection cycle, this yields an estimated ~2.78 mutations per cycle.

**1.3 Contribution of *E. coli* replication and recombination to YFP-MutL foci counts.** We observed ~3 times more YFP-MutL foci than predicted from the lambda replication alone, suggesting additional contributions from replication errors on the host genome. Although initiation of new replication forks on the *E. coli* chromosome is blocked following lambda induction, elongation continues on forks that were already active at the time of infection (Wold et al. 1982; Sergueev et al. 2002).

The amount of *E. coli* DNA synthesized during lambda infection depends on the cell cycle stage at the time of infection, i.e., the number of active replication forks. Under our growth conditions (LB medium, doubling time ~26 minutes), *E. coli* cells typically contain 6 forks between 0–6 minutes of age, 4 forks between 6–18 minutes, and 12 forks between 18–26 minutes (Zaritsky et al. 2011). Given the ~40-minute duration of the lytic cycle, we estimate that between 6 Mb (for cells with 4 forks, infected at age 6 min) and 22 Mb (for cells with 12 forks infected at age 18 minutes) of *E. coli* DNA can be replicated during lambda infection. This represents 17–43% of total DNA synthesis per infection cycle.

Replication errors on the *E. coli* genome may be partially or fully detected as YFP-MutL foci, depending on the availability of MutS and MutL proteins and the efficiency of MMR on the host genome during lambda infection. Assuming sufficient levels of MutS and MutL and efficient repair of replication errors, approximately 65% of these events would be detectable under our imaging conditions (1-minute intervals), due to the short average lifetime of MutL foci on repairable errors (~40 seconds; see Enrico Bena et al., 2024). Under these conditions, replication errors on the host genome are estimated to contribute 11–28% of all observed foci.

Conversely, if MMR is impaired during infection (i.e., MutL foci persisting for ≥25 minutes), nearly all replication errors could be detected, with host-derived errors contributing up to 17–43% of total foci. In this extreme scenario, in some cells up to half of the observed YFP-MutL foci could originate from replication errors on the *E. coli* genome. During the lytic cycle, additional mismatches may arise from recombination between lambda and the defective prophages Dlp12 and Qin, which reside on the *E. coli* genome and share regions of high sequence homology with lambda (see MA-WGS section). Recombination occurs primarily with the defective prophage Dlp12, as shown by mismatch clusters in 11 of the 12 MA lines. These recombination-associated mismatches should, in principle, be detectable as YFP-MutL foci. However, we observed no change in the rate of YFP-MutL foci per infection cycle upon deletion of the region of Dlp12 homologous to lambda (Table S10), suggesting that such events are either rare or too transient to be reliably captured under our imaging conditions.

**1.4 Accuracy of our estimates of lambda mutation rates.** We employed three different methods to estimate the mutation rate of lambda: the FA, performed during a single-burst experiment, and DS and MA-WGS, both of which detect mutations arising over multiple rounds of infection. Because these approaches differ in both experimental design and what they measure, the mutation rate is calculated differently for each. In the fluctuation assay, the per-replication mutation rate is obtained by dividing the inferred number of mutations by the estimated number of phage DNA replications. In contrast, DS and MA-WGS estimate the mutation rate by dividing the total number of detected mutations by the estimated number of phage DNA doublings. The accuracy of each method's mutation rate estimate depends on how accurately we estimate or measure both the number of mutations and the number of DNA replications or doublings. Each method introduces specific biases in these estimates, as discussed below.

We estimate the total number of DNA replications and doublings from the total phage count (at the end of each experiment). The number of DNA replications is straightforward to estimate as the total number of phages produced minus one. In contrast, estimating the number of DNA doublings requires assumptions about both the replication mechanism and infection dynamics. We assume that lambda replicates exclusively via bidirectional replication and that each progeny phage infects a new bacterium. However, in reality, lambda replication cycle is more complex, consisting of 5–6 rounds of bidirectional replication within a single infection, followed by one round of rolling-circle replication (Taylor and Węgrzyn 1995). Additionally, not all phages successfully initiate new infections, particularly in later cycles. As a result, our calculation likely underestimates the true number of DNA doublings, leading to an overestimation of the mutation rate from DS and MA-WGS datasets (Supplementary Information 1.5).

In DS and MA-WGS, mutations are directly detected and quantified through sequencing. In contrast, the FA infers the number of mutations based on the observed number of mutant phages. This inference is typically performed using the Ma-Sandri-Sarkar (MSS) maximum likelihood estimator, which assumes exponential growth, a condition that does not accurately reflect the replication dynamics of lambda. To overcome this limitation, we also applied the $P_0$ method, which does not require assumptions about the mode of replication. Instead, it estimates the number of mutation events based solely on whether any mutation occurred in a culture.

Given these considerations, we regard our FA-based estimate derived using the $P_0$ method and the $cII_{G183A}$ phage as the most accurate estimate of the lambda mutation rate, as it is likely the least affected by assumptions about phage growth dynamics. Yet, the estimate obtained from FA assay is based on mutations selected within a small genomic region and may not accurately reflect genome-wide mutation rate. Studies in bacteria and yeast have shown that the chromosomal location of the reporter gene can lead to several-fold variation in observed mutation rates (Foster et al. 2013; Long et al. 2014; Dillon et al. 2015, 2017, 2018). We have shown here the absence of important regional variation in mutation rate on lambda chromosome, yet whether small regional variation occurs in lambda remains to be determined with a larger collection of mutants.

**1.5 Estimating the extent of mutation rate overestimation in MA experiment.** The number of DNA doublings per line was estimated from 74 measurements of the number of PFU per lysis plaque, whose mean is equal to 5.9 ± 0.3 x$10^7$ PFUs. Assuming exponential growth, this corresponds to $Log_2$(6 x$10^7$) = 25.8 genome replications. Assuming that about 240 particles are produced per cell and per lytic cycle, this roughly corresponds to 4 successive lytic cycles. Given the low adsorption rate of the lambda *stf*- phage used (Shao and Wang 2008), we can approximate that it takes more than 1.9 hour for the first infectious lytic cycle to start and more than 20 minutes between the first and the second cycle. These numbers are underestimation as it uses a phage adsorption rate in liquid, that must be lower in top agar due to lower diffusion than in water. Altogether, if we add 2.5 hours for infection and 4 hours in lytic cycles it makes 6.5 hours, yet it takes 8 hours to obtain 6 x$10^7$ phages, which could correspond to a $5^{th}$ lytic cycle necessary to compensate phage particle lost due to absorption on cell debris or on already infected bacteria. Assuming 5 lytic cycles would correspond to 33 doubling instead of 26, a 27% increase, that would lead to a 30% overestimation of mutation rate. Similarly, assuming 15 % of replication in linear mode (5-6 bidirectional replications, followed by a single round of rolling circle for matured DNA) would lead to 15% overestimation of the mutation rate. Combining both assumptions would lead to a 50% overestimation of the rate and thus and a 1/$3^{rd}$ decrease of the real mutation rate compared to the estimated one.

### Supplementary figures

**Figure S1**

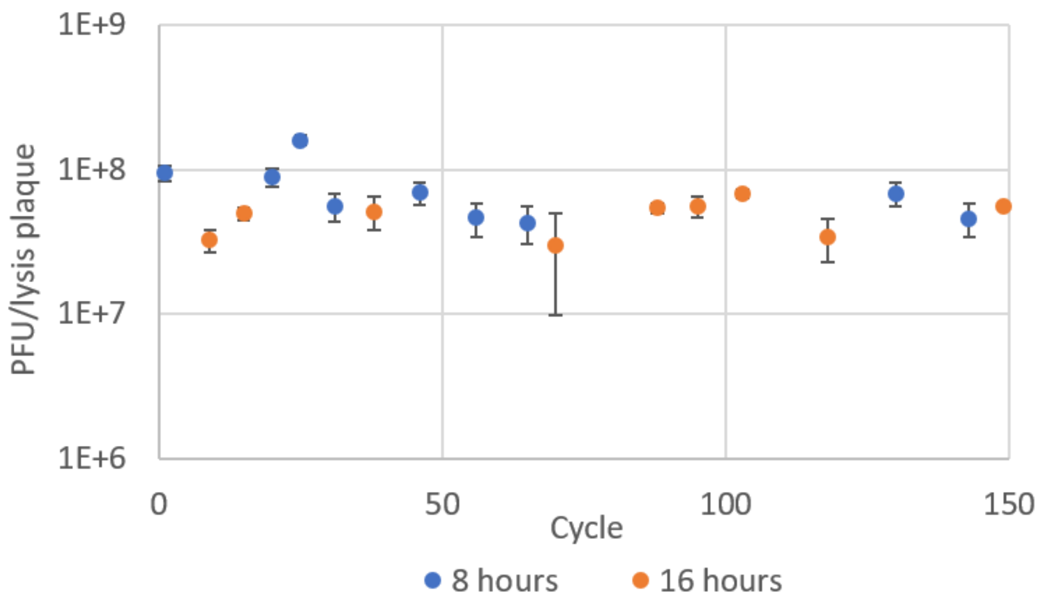

**Figure S1. No evolution of the number of PFU per plaque with time.** Each dot represents the mean of four lines +/- sem. No evolution with time is observed, indicating the absence of selection of strongly beneficial or deleterious mutations in these four lines. Furthermore, no difference is observed between 8- and 16-hours incubation time.

**Figure S2**

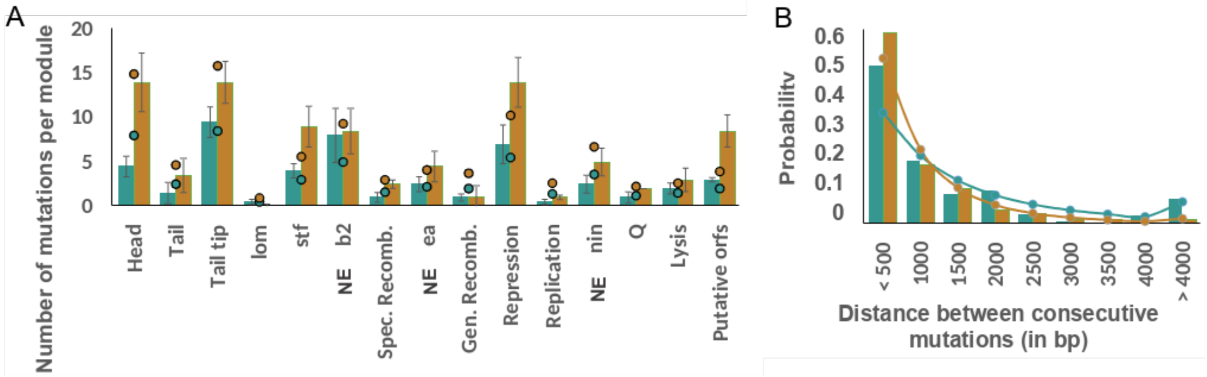

**Figure S2. No mutation hotspots are observed in the DS experiment. A)** Number of mutations per transcription module (bar plot). The dots represent the number of mutations expected per module if mutations were regularly distributed. The modules' name and limits were determined from a review (Rajagopala et al. 2011). NE = non-essential. **B)** The frequency of the distance between consecutive mutations per 500 bp interval is plotted as bar plot for the samples pooled per strain. The probability distribution of the distances under the hypothesis of random distribution of mutations (exponential law) is plotted as a dotted line. blue = WT; orange = *mutS*.

**Figure S3**

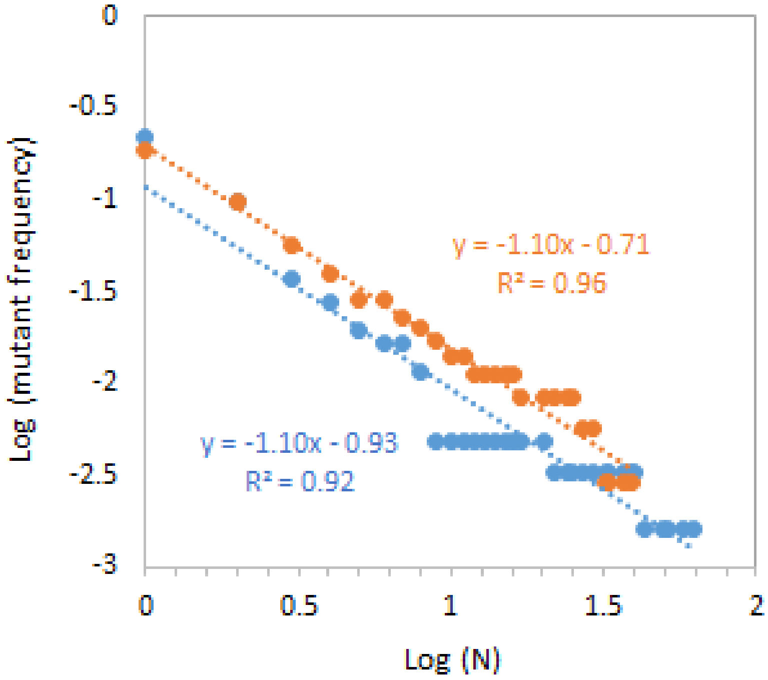

**Figure S3. The distribution of the number of lambda *cII* mutants per sample follows a Luria-Delbrück distribution.** The $\log_{10}$ of the number of cultures with x or more mutants is plotted against the $\log_{10}$ of the number of mutants per culture. The dotted lines correspond to linear trendlines, with the corresponding equation and coefficient of determination ($R^2$) indicated. Blue : lambda multiplied on a WT *E. coli* strain, orange : lambda multiplied on a *mutS E. coli* strain.

**Figure S4**

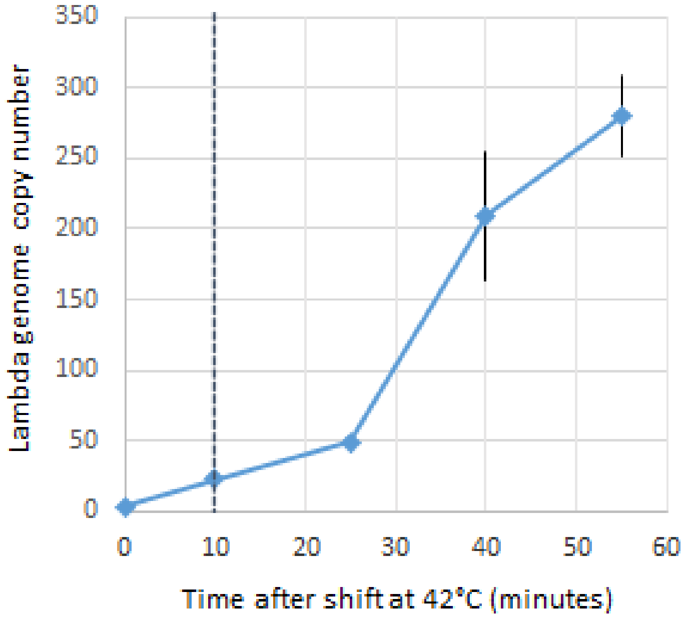

**Figure S4: Evolution with time of the mean number of lambda genomes per cell following prophage induction.** From 0 to 10 minutes, the lysogenic culture is incubated at 42°C. Then, cells are incubated at 37°C until 60 min. The lambda genome copy number is determined by semi-quantitative PCR, relatively to the number of *E. coli* genome copy number, and hypothesizing a mean number of 3 *E. coli* genomes per cell.

**Figure S5**

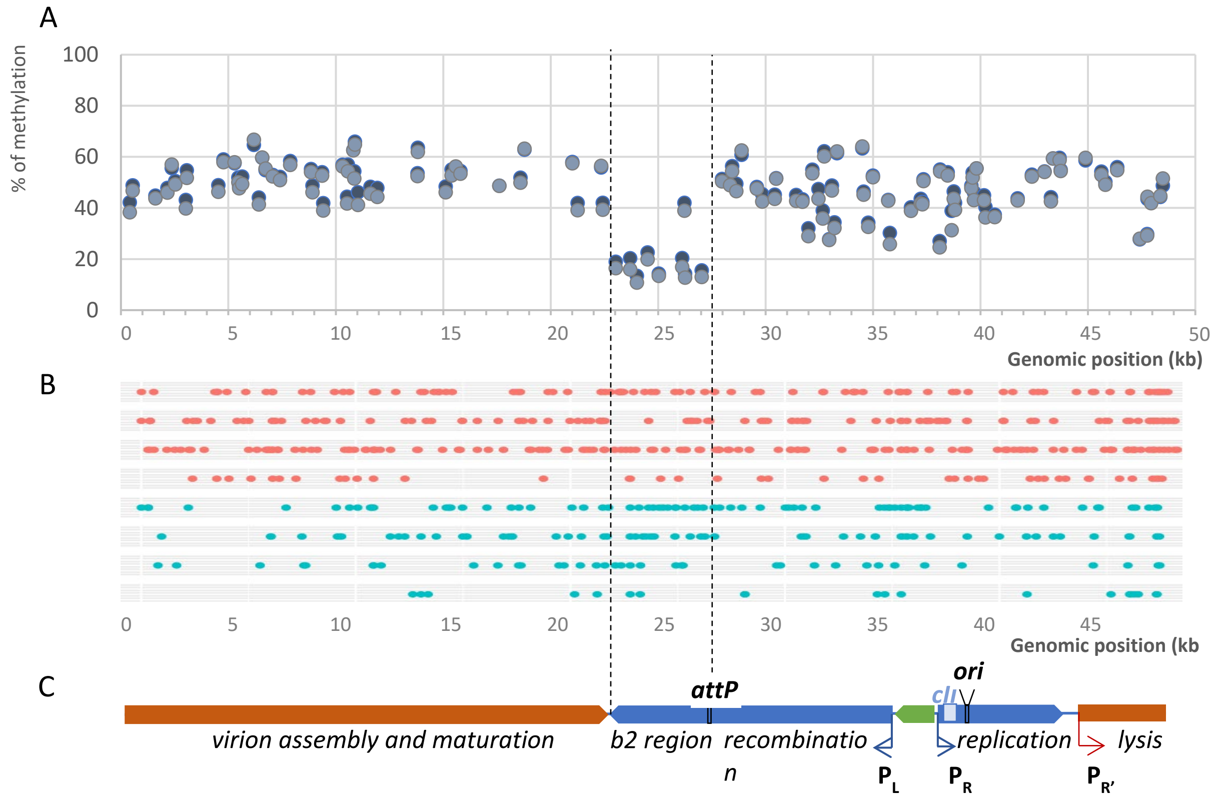

**Figure S5. lambda GATC undermethylation does not impact the distribution of mutations**. **A)** Methylation level of adenines at each GATC site in lambda genome. Results of two independent replicates are shown, depicted in blue and grey respectively. **B)** Positions on the lambda genome of the mutations determined in the DS experiments. The 4 replicates per condition are plotted. Red = *mutS*; Blue = WT. **C)** Representation of the main transcription units of lambda genome. Green = lysogeny control region; Blue = Early genes; Red = Late genes. The vertical dotted lines delimit the region comprised between *attB* and the terminus of $P_{R'}$ operon, the most expressed during the lytic cycle. We can see it delimits a region of lower levels of methylation in A), probably due to conflict between transcription and methylation.

**Figure S6**

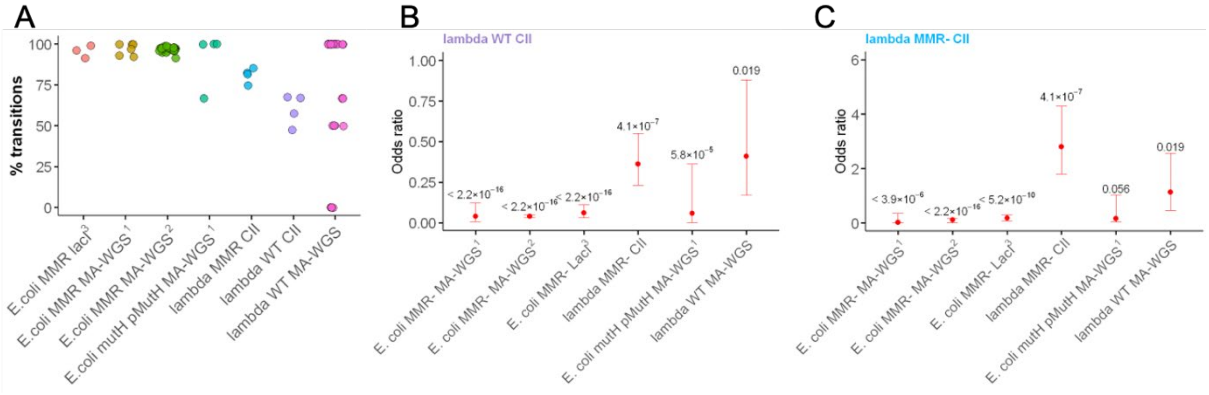

**Figure S6. Lambda and E. coli chromosomes display a different mutation profile in MMR deficient strains. A)** Percentage of transitions in lambda *vs* MMR deficient *E. coli.* Mutations were obtained by selection for *cII* (lambda) or *lacI* inactivation (*E. coli*) or by MA-WGS (lambda and *E. coli*). Data for *E. coli* spectra are from: [1]Enrico Bena et al., 2024, *Nature Communications*; [2]Niccum et al., 2018, *Genetics*; [3]Schaaper et al., 1987, *PNAS*. Each colored dot represents an independent experiment. **B)** Statistical comparisons of transversion-to-transition ratios between lambda grown on WT or **C)** MMR deficient *E. coli* and the conditions indicated on the x-axis. Red dots indicate odds ratios from Fisher's exact test; error bars represent 95% confidence intervals, p-values shown above dots indicate statistical significance.

### Supplementary tables

**Table S1: Mutation frequencies per base sequenced obtained in the DS experiment.** *Frequencies for WT and *mutS* are shown after subtracting the background mutation frequency observed in pUC19. N, number; BPS, base pair substitutions

| Genotype | Replicate | N BPS | N indels | N bases sequenced | BPS frequency ($x10^{-7}$)* | Indels frequency ($x10^{-7}$)* |
|---|---|---|---|---|---|---|
| pUC19 | 1 | 6 | 2 | 41 883 220 | 1.43 | 0.48 |
| WT | 1 | 64 | 9 | 99 768 081 | 4.98 | 0.42 |
| | 2 | 48 | 11 | 75 730 852 | 4.91 | 0.98 |
| | 3 | 31 | 6 | 42 984 793 | 5.78 | 0.92 |
| | 4 | 16 | 2 | 32 739 555 | 3.46 | 0.74 |
| *mutS* | 1 | 70 | 19 | 61 762 461 | 9.87 | 2.59 |
| | 2 | 94 | 6 | 68 394 037 | 12.27 | 0.4 |
| | 3 | 109 | 24 | 100 246 812 | 9.47 | 1.92 |
| | 4 | 36 | 5 | 30 366 202 | 10.47 | 1.17 |

**Table S2: Mutation rates per locus per replication obtained in the Fluctuation assay.** Missing values for the $P_0$ estimator are due to the absence of the null class in the distribution of mutants per culture for some samples.

| Fluctuation assay parameters | | | Estimator (mutation rate per locus $x10^{-6}$) | |
|---|---|---|---|---|
| Host genotype | Phage genotype | Experiment | $P_0$ | Salvador |
| WT | WT | 1 | 5.84 | 4.73 |
| WT | WT | 2 | 4.84 | 4.57 |
| WT | WT | 3 | 5.57 | 4.61 |
| WT | WT | 4 | 4.42 | 3.50 |
| WT | WT | 5 | 5.60 | 4.62 |
| WT | WT | 6 | 4.29 | 3.17 |
| WT | WT | 7 | 3.05 | 2.75 |
| WT | WT | 8 | 6.98 | 5.15 |
| WT | WT | 9 | 5.40 | 4.58 |
| WT | WT | 10 | 6.46 | 4.81 |
| WT | WT | 11 | 11.00 | 7.20 |
| WT | WT | 12 | 6.40 | 6.20 |
| WT | WT | 13 | 7.50 | 7.60 |
| WT | WT | 14 | 6.10 | 4.30 |
| WT | WT | 15 | 5.30 | 4.10 |
| WT | WT | 16 | 4.90 | 3.50 |
| WT | WT | 17 | 5.10 | 4.10 |
| WT | WT | 18 | 13.00 | 9.30 |
| *mutS* | WT | 1 | 13.09 | 7.60 |
| *mutS* | WT | 2 | 38.46 | 42.70 |

| | | | | |
|---|---|---|---|---|
| *mutS* | WT | 3 | 73.85 | 59.80 |
| *mutS* | WT | 4 | 64.61 | 54.90 |
| *mutS* | WT | 5 | 96.11 | 102.00 |
| *mutS* | WT | 6 | 60.09 | 44.90 |
| *mutS* | WT | 7 | - | 32.70 |
| *dam* | WT | 1 | - | 15.90 |
| *dam* | WT | 2 | 17.20 | 15.70 |
| *dam* | WT | 3 | 13.20 | 10.00 |
| *dam* | WT | 4 | 5.60 | 4.20 |
| *dam* | WT | 5 | - | 30.50 |
| *dam* | WT | 6 | 25.10 | 17.70 |
| WT | G183A | 1 | 0.92 | 0.55 |
| WT | G183A | 2 | - | 0.57 |
| WT | G183A | 3 | - | 0.67 |
| WT | G183A | 4 | - | 0.95 |
| WT | G183A | 5 | - | 1.63 |
| *mutS* | G183A | 1 | 16.0 | 8.70 |
| *mutS* | G183A | 2 | 17.0 | 11.00 |
| *mutS* | G183A | 3 | 11.0 | 8.10 |
| *mutS* | G183A | 4 | 8.8 | 5.70 |
| *mutS* | G183A | 5 | 4.9 | 4.10 |
| *mutS* | G183A | 6 | 11.0 | 6.80 |
| *mutS* | G183A | 7 | 7.7 | 6.20 |

**Table S3: Distribution of CII inactivating mutations.** Positions and BPS observed in *cII* Sanger sequencing. Stop codons are indicated by their name (UGA = opal, UAG = amber, UAA = ochre). Under parentheses are the BPS that were deleted from the analysis, as these result from the 6G mutagenesis hotspot.

| Site | AA change | bp change | WT | MMR- |
|---|---|---|---|---|
| 2 | M1I | T>C | 0 | 1 |
| 14 | N5S | A>G | 1 | 0 |
| 15 | N5K | C>A | 1 | 0 |
| 25 | E9amber | G>T | 0 | 1 |
| 28 | A10T | G>A | 2 | 6 |
| 28 | A10P | G>C | 1 | 0 |
| 29 | A10D | C>A | 9 | 4 |
| 29 | A10V | C>T | 2 | 5 |
| 31 | L11V | C>G | 1 | 0 |
| 31 | L11L | C>T | 0 | 1 |
| 32 | L11P | T>C | 2 | 2 |
| 34 | R12opal | C>T | 1 | 1 |
| 35 | R12Q | G>A | 1 | 0 |
| 35 | R12P | G>C | 0 | 1 |
| 37 | I13F | A>T | 1 | 0 |

| | | | | |
|---|---|---|---|---|
| 38 | I13T | T>C | 0 | 1 |
| 40 | E14K | G>A | 3 | 0 |
| 40 | E14amber | G>T | 1 | 1 |
| 49 | L17M | T>A | 2 | 0 |
| 50 | L17S | T>C | 3 | 3 |
| 51 | L17F | G>T | 6 | 1 |
| 55 | N19D | A>G | 0 | 1 |
| 56 | N19S | A>G | 1 | 2 |
| 57 | N19K | C>A | 4 | 0 |
| 58 | K20E | A>G | 1 | 0 |
| 58 | K20ochre | A>T | 1 | 0 |
| 60 | K20N | A>C | 0 | 1 |
| 61 | I21V | A>G | 3 | 0 |
| 62 | I21S | T>G | 1 | 0 |
| 65 | A22E | C>A | 3 | 1 |
| 65 | A22G | C>G | 0 | 1 |
| 65 | A22V | C>T | 1 | 0 |
| 68 | M23T | T>C | 1 | 3 |
| 71 | L24P | T>C | 1 | 4 |
| 74 | G25E | G>A | 1 | 2 |
| 74 | G25A | G>C | 0 | 2 |
| 74 | G25V | G>T | 3 | 0 |
| 76 | T26P | A>C | 0 | 1 |
| 76 | T26A | A>G | 4 | 1 |
| 79 | E27amber | G>T | 0 | 1 |
| 81 | E27E | G>A | 1 | 0 |
| 82 | K28E | A>G | 3 | 0 |
| 85 | T29P | A>C | 0 | 1 |
| 86 | T29K | C>A | 2 | 0 |
| 87 | T29T | A>T | 1 | 0 |
| 88 | A30T | G>A | 3 | 1 |
| 88 | A30S | G>T | 1 | 0 |
| 89 | A30D | C>A | 3 | 1 |
| 89 | A30V | C>T | 3 | 2 |
| 91 | E31ochre | G>T | 1 | 0 |
| 94 | A32T | G>A | 2 | 0 |
| 95 | A32D | C>A | 2 | 0 |
| 98 | V33A | T>C | 0 | 9 |
| 100 | G34S | G>A | 2 | 1 |
| 101 | G34D | G>A | 1 | 2 |
| 101 | G34A | G>C | 1 | 0 |
| 101 | G34V | G>T | 1 | 3 |
| 103 | V35I | G>A | 2 | 0 |
| 103 | V35F | G>T | 0 | 1 |
| 104 | V35A | T>C | 1 | 5 |
| 104 | V35G | T>G | 0 | 2 |

| | | | | |
|---|---|---|---|---|
| 107 | D36G | A>G | 0 | 3 |
| 109 | K37E | A>G | 1 | 0 |
| 111 | K37N | G>T | 0 | 1 |
| 112 | S38P | T>C | 1 | 6 |
| 113 | S38L | C>T | 0 | 1 |
| 115 | Q39K | C>A | 0 | 1 |
| 115 | Q39E | C>G | 1 | 0 |
| 115 | Q39amber | C>T | 1 | 0 |
| 116 | Q39L | A>T | 1 | 0 |
| 117 | Q39H | G>T | 1 | 0 |
| 118 | I40L | A>C | 1 | 0 |
| 118 | I40V | A>G | 0 | 2 |
| 119 | I40N | T>A | 1 | 1 |
| 119 | I40T | T>C | 4 | 2 |
| 119 | I40S | T>G | 3 | 0 |
| 121 | S41R | A>C | 1 | 0 |
| 121 | S41G | A>G | 6 | 5 |
| 122 | S41I | G>T | 1 | 0 |
| 122 | S41N | G>A | 1 | 0 |
| 123 | S41R | C>A | 3 | 0 |
| 124 | R42G | A>G | 3 | 2 |
| 124 | R42W | A>T | 1 | 0 |
| 126 | R42S | G>T | 2 | 0 |
| 127 | W43R | T>A | 4 | 0 |
| 127 | W43R | T>C | 12 | 24 |
| 127 | W43G | T>G | 1 | 2 |
| 128 | W43amber | G>A | 1 | 0 |
| 128 | W43L | G>T | 1 | 1 |
| 129 | W43opal | G>A | 2 | 4 |
| 129 | W43C | G>T | 1 | 1 |
| 130 | K44Q | A>C | 1 | 0 |
| 130 | K44E | A>G | 2 | 3 |
| 131 | K44R | A>G | 1 | 0 |
| 132 | K44N | G>T | 3 | 0 |
| 133 | R45G | A>G | 0 | 2 |
| 133 | R45W | A>T | 0 | 1 |
| 136 | D46Y | G>T | 0 | 1 |
| 137 | D46G | A>G | 5 | 1 |
| 139 | W47R | T>C | 1 | 0 |
| 140 | W47amber | G>A | 2 | 3 |
| 141 | W47opal | G>A | 4 | 3 |
| 141 | W47C | G>C | 1 | 0 |
| 141 | W47C | G>T | 1 | 0 |
| 143 | I48T | T>C | 3 | 4 |
| 145 | P49T | C>A | 2 | 0 |
| 145 | P49S | C>T | 1 | 1 |

| | | | | |
|---|---|---|---|---|
| 148 | K50Q | A>C | 1 | 0 |
| 148 | K50E | A>G | 2 | 1 |
| 149 | K50T | A>C | 1 | 0 |
| 149 | K50R | A>G | 2 | 0 |
| 150 | K50N | G>T | 4 | 0 |
| 152 | F51S | T>C | 0 | 4 |
| 152 | F51C | T>G | 2 | 0 |
| 161 | L54P | T>C | 5 | 8 |
| 163 | L55I | C>A | 4 | 2 |
| 163 | L55F | C>T | 2 | 3 |
| 164 | L55P | T>C | 1 | 0 |
| 166 | A56T | G>A | 2 | 0 |
| 166 | A56S | G>T | 1 | 0 |
| 167 | A56V | C>T | 0 | 1 |
| 169 | V57F | G>T | 3 | 0 |
| 173 | L58P | T>C | 0 | 2 |
| 175 | E59ochre | G>T | 1 | 0 |
| 176 | E59C | A>C | 1 | 1 |
| 177 | E59E | A>G | 0 | 1 |
| 178 | W60R | T>C | 2 | 2 |
| 179 | W60amber | G>A | 1 | 0 |
| 179 | W60S | G>C | 1 | 0 |
| 180 | W60opal | G>A | 5 | 5 |
| 182 | G61V | GG>TC | 1 | 0 |
| 185 | V62A | T>C | 0 | 1 |
| 185 | V62G | T>G | 0 (40) | 0 (4) |
| 187 | V63F | G>T | 1 | 0 |
| 191 | D64V | A>C | 1 | 1 |
| 191 | D64G | A>G | 7 | 6 |
| 193 | D65N | G>A | 1 | 0 |
| 194 | D65G | A>G | 3 | 1 |
| 196 | D66N | G>A | 1 | 0 |
| 196 | D66H | G>C | 1 | 0 |
| 196 | D66Y | G>T | 2 | 0 |
| 197 | D66A | A>C | 0 | 1 |
| 197 | D66G | A>G | 5 | 6 |
| 200 | M67T | T>C | 0 | 2 |
| 202 | A68S | G>T | 1 | 0 |
| 205 | R69opal | C>T | 0 | 1 |
| 209 | L70S | T>C | 1 | 0 |
| 210 | L70F | G>C | 1 | 0 |
| 210 | L70F | G>T | 1 | 0 |
| 211 | A71T | G>A | 3 | 0 |
| 212 | A71E | C>A | 1 | 1 |
| 212 | A71V | C>T | 6 | 1 |
| 214 | R72opal | C>T | 3 | 3 |

| | | | | |
|---|---|---|---|---|
| 217 | Q73ochre | C>T | 0 | 1 |
| 220 | V74F | G>T | 1 | 2 |
| 221 | V74A | T>C | 2 | 3 |
| 223 | A76T | G>A | 3 | 0 |
| 224 | A76V | C>T | 1 | 0 |
| 226 | A76T | G>A | 0 | 1 |
| 226 | A76P | G>C | 1 | 0 |
| 230 | I77T | T>C | 2 | 0 |
| 230 | I77M | T>G | 0 | 1 |
| 231 | I77S | T>G | 4 | 1 |
| 233 | L78P | T>C | 1 | 2 |
| 233 | L78R | T>G | 0 | 1 |
| 235 | T79P | A>C | 0 | 1 |
| 292 | opal98R | T>A | 1 | 0 |
| 293 | opal98L | G>T | 1 | 0 |

**Table S4: Transitions vs. transversions in *E. coli* and lambda, as determined by fluctuation assays (*lacI*, *cII*) and MA-WGS.** (Data used to generate Figure 4A). [1]Enrico Bena et al., 2024, *Nature Com.*; [2]Niccum et al., 2018, *Genetics*; [3]Schaaper et al., 1987, *PNAS*.

| Organism | lambda | *E. coli* | *E. coli* | *E. coli* | lambda | lambda | *E. coli* |
|---|---|---|---|---|---|---|---|
| Host background | WT | MMR- | MMR- | MMR- | WT | MMR- | *mutH pMutH* |
| Experiment Type | MA-WGS | MA-WGS[1] | MA-WGS[2] | LacI-mutants[3] | CII-mutants | CII-mutants | MA-WGS[1] |
| Transitions | 34 | 106 | 29,234 | 350 | 158 | 173 | 24 |
| Transversions | 10 | 3 | 827 | 15 | 115 | 45 | 1 |
| Total | 44 | 109 | 30,061 | 365 | 273 | 218 | 25 |
| % Transitions | 77 | 97 | 97 | 96 | 58 | 79 | 96 |

**Table S5: Mutation spectrum of BPS in *E. coli* inactivated for MMR or showing partial MMR activity, as determined by MA-WGS** (Enrico Bena et al. 2024)

| Strain | *mutH* | *mutH* | *mutH* | *mutH* | *mutH* | *mutH* | *mutH pMutH* | *mutH pMutH* | *mutH pMutH* | *mutH pMutH* |
|---|---|---|---|---|---|---|---|---|---|---|
| Experiment | 1 | 2 | 3 | 4 | 5 | 6 | 1 | 2 | 3 | 4 |
| G>A | 1 | 2 | 2 | 4 | 0 | 3 | 0 | 1 | 0 | 2 |
| C>T | 2 | 2 | 3 | 4 | 1 | 7 | 1 | 0 | 2 | 0 |
| A>G | 10 | 5 | 1 | 5 | 6 | 9 | 0 | 3 | 4 | 2 |
| T>C | 4 | 4 | 6 | 6 | 7 | 12 | 1 | 2 | 5 | 1 |
| G>T | 0 | 0 | 0 | 0 | 0 | 1 | 0 | 0 | 0 | 0 |
| C>A | 0 | 0 | 0 | 0 | 0 | 0 | 0 | 0 | 0 | 0 |
| T>G | 0 | 1 | 0 | 0 | 0 | 0 | 0 | 0 | 0 | 0 |
| A>C | 0 | 0 | 0 | 0 | 0 | 0 | 1 | 0 | 0 | 0 |
| G>C | 0 | 0 | 0 | 0 | 0 | 0 | 0 | 0 | 0 | 0 |

| | | | | | | | | | | |
|---|---|---|---|---|---|---|---|---|---|---|
| C>G | 0 | 0 | 1 | 0 | 0 | 0 | 0 | 0 | 0 | 0 |
| A>T | 0 | 0 | 0 | 0 | 0 | 0 | 0 | 0 | 0 | 0 |
| T>A | 0 | 0 | 0 | 0 | 0 | 0 | 0 | 0 | 0 | 0 |
| Transitions | 17 | 13 | 12 | 19 | 14 | 31 | 2 | 6 | 11 | 5 |
| Transversions | 0 | 1 | 1 | 0 | 0 | 1 | 1 | 0 | 0 | 0 |
| Total | 17 | 14 | 13 | 19 | 14 | 32 | 3 | 6 | 11 | 5 |

**Table S6: Odd ratios and 95% confidence intervals (in parenthesis) as obtained by Fisher Exact test on the mutation rates.** [1](Niccum et al. 2018); [2](Enrico Bena et al. 2024); [3](Schaaper and Dunn 1987)

| | lambda WT MA-WGS | *E. coli* MMR- MA-WGS[1] | *E. coli* MMR- MA-WGS[2] | *E. coli* MMR- *lacI*[3] | lambda WT CII | lambda MMR- CII | *E. coli* *mutH* *pmutH* MA-WGS[2] |
|---|---|---|---|---|---|---|---|
| lambda WT MA-WGS | 1 (0.32-3.1) | 0.098 (0.016-0.41) | 0.096 (0.046-0.22) | 0.15 (0.057-0.4) | 2.4 (1.1-5.8) | 0.89 (0.39-2.2) | 0.15 (0.003-1.1) |
| *E. coli* MMR- MA-WGS[1] | 10 (2.4-61) | 1 (0.13-7.6) | 1 (0.33-4.9) | 1.5 (0.42-8.3) | 26 (8.2-130) | 9.12 (2.8-47) | 1.5 (0.027-19) |
| *E. coli* MMR- MA-WGS[2] | 10 (4.6-22) | 1 (0.20-3.015) | 1 (0.9-1.1) | 1.5 (0.83-2.6) | 26 (20-33) | 9.2 (6.4-13) | 1.5 (0.036-9.057) |
| *E. coli* MMR- *lacI*[3] | 6.8 (2.5-18) | 0.66 (0.12-2.4) | 0.66 (0.39-1.2) | 1 (0.45-2.2) | 17 (9.5-32) | 6.05 (3.2-12) | 0.97 (0.022-6.9) |
| lambda WT CII | 0.41 (0.17-0.88 | 0.039 (0.008-0.12) | 0.039 (0.03-0.05) | 0.059 (0.031-0.11) | 1 (0.7-1.4) | 0.36 (0.23-0.55) | 0.057 (0.001-0.36) |
| lambda MMR- CII | 1.13 (0.46-2.56) | 0.11 (0.021-0.35 | 0.11 (0.078-0.16) | 0.17 (0.083-0.31) | 2.8 (1.8-4.3) | 1 (0.61-1.6) | 0.16 (0.038-1.04) |
| *E. coli* *mutH* *pmutH* MA-WGS[2] | 6.9 (0.88-320) | 0.68 (0.052-37) | 0.68 (0.11-28) | 1 (0.15-45) | 17 (2.8-720) | 6.2 (0.96-260) | 1 (0.012-81) |

**Table S7: p-values as obtained by Fisher Exact test on the mutation rates.** [1](Niccum et al. 2018); [2](Enrico Bena et al. 2024); [3](Schaaper and Dunn 1987)

| | lambda WT MA-WGS | *E. coli* MMR- MA-WGS[1] | *E. coli* MMR- MA-WGS[2] | *E. coli* MMR- *lacI*[3] | lambda WT CII | lambda MMR- CII | *E. coli* *mutH* *pmutH* MA-WGS[2] |
|---|---|---|---|---|---|---|---|
| lambda WT MA-WGS | 1 | 0.00024 | $2.8 \times 10^{-7}$ | $7.4 \times 10^{-5}$ | 0.019 | 0.84 | 0.047 |
| *E. coli* MMR- MA-WGS[1] | 0.00024 | 1 | 1 | 0.78 | $< 2.2 \times 10^{-16}$ | $3.9 \times 10^{-6}$ | 0.57 |
| *E. coli* MMR- MA-WGS[2] | $2.8 \times 10^{-7}$ | 1 | 1 | 0.14 | $< 2.2 \times 10^{-16}$ | $< 2.2 \times 10^{-16}$ | 0.5 |
| *E. coli* MMR- *lacI*[3] | $7.4 \times 10^{-5}$ | 0.78 | 0.14 | 1 | $< 2.2 \times 10^{-16}$ | $5.2 \times 10^{-10}$ | 1 |
| lambda WT CII | 0.019 | $< 2.2 \times 10^{-16}$ | $< 2.2 \times 10^{-16}$ | $< 2.2 \times 10^{-16}$ | 1 | $4.1 \times 10^{-7}$ | $5.8 \times 10^{-5}$ |
| lambda MMR- | 0.84 | $3.9 \times 10^{-6}$ | $< 2.2 \times 10^{-16}$ | $5.2 \times 10^{-10}$ | $4.1 \times 10^{-7}$ | 1 | 0.056 |

| | | | | | | | |
|---|---|---|---|---|---|---|---|
| CII | | | | | | | |
| *E. coli mutH pmutH* MA-WGS[2] | 0.047 | 0.57 | 0.50 | 1 | $5.8 \times 10^{-5}$ | 0.056 | 1 |

**Table S8: List of bacterial strains, phages, and plasmids used in this study**

| *E. coli* strains/plasmids | Genotype | Source |
|---|---|---|
| MG1655 6300 | WT | GSC |
| MG1655 | WT | Ivan Matic's lab collection |
| DH10B | F- endA1 recA1 galE15 galK16 nupG rpsL ΔlacX74 Φ80lacZΔM15 araD139 Δ(ara,leu)7697 mcrA Δ(mrr-hsdRMS-mcrBC) lambda- | (Grant et al. 1990) |
| GSY5902 | AB1157 *ΔrecA306 srl::Tn10* (*miniF recA*) | (Martinsohn et al. 2008) |
| ME120 | MG1655 *plac-yfp-mutL-cat mutL218::Tn10* | (Elez et al. 2012) |
| ME120R | ME120 *attTn7::pRNA1-tdCherry* | This study |
| 63ME120R | MG1655 6300 *lacZ::yfp-mutL-frt, mutL::frt, attTn7::pRNA1-tdCherry* | (Robert et al. 2018) |
| 63ME120RL | 63ME120R lambda *Ind1* $cI_{857}$ | This study |
| L6 | 63ME120R HomDlp12::FRT (*rzpD-tfaX*) *lambda*$_{cI857}$ | This study |
| L12 | 63ME120R *mutS::SmR* lambda *Ind1* $cI_{857}$ | This study |
| JL10 | MG1655 *stfR*::kan | This study |
| GSY5902 | AB1157 *ΔrecA306 srl::*Tn10 *(miniF recA)* | Martinsohn et al., Plos Genet. 2008 |
| MD19 | MG1655 tfaQ::*cat* | (De Paepe et al. 2014) |
| MD97 | MG1655 *polB::kan dinB::frt umuCD::cat* | This study |
| SL311 | MG1655 6300 *lac-3350 galK2 galT22 rpsL179 hflB29 zgj25::Tn10 hflA::Tn5* | (Herman et al. 1993) |
| *ME14* | MG1655 6300 *mutS::cat* | This study |
| MVEC249 | MG1655 6300 *mutS::SmR* | This study |
| MVEC318 | MG1655 6300 lambda *Ind1* $cI_{857}$ $cII_{G183A}$ | This study |
| MVEC319 | MG1655 6300 lambda *Ind1* $cI_{857}$ $cII_{G183A}$ | This study |
| MD114 | MG1655 lambda *Ind1* $cI_{857}$ | This study |
| MVEC235 | MG1655 6300 lambda *Ind1* $cI_{857}$ | This study |
| MD135 | MG1655 *dam::cat* | Ivan Matic's lab collection |
| MVEC197 | MG1655 6300 *pLtetO1-dam* | (Enrico Bena et al. 2024) |
| MGZ1X | MG1655 Z1 ($P_{N25}$*-tetR* $P_{lacI}^{q}$*-lacI araC*) | (Cox et al. 2007) |
| MVEC232 | MD135 lambda *Ind1* $cI_{857}$ | This study |
| MVEC276 | MVEC235 *recA306 srl::Tn10-tetR* | This study |
| MVEC333 | MG1655 6300 *lacZ::yfp-mutL-frt, mutL::frt, attTn7::pRNA1-tdCherry ΔfliC::FRT* | This study |
| MVEC337 | MG1655 6300 *lacZ::Plac-yfp-mutL; mutL::FRT ; fliC::FRT ; lcI857 bor::cm-21tetO; pACYC177-pFtski-tetR-mCherry* | This study |
| MD301 | MGZ1X *pLTetO1-dam* | This study |
| Plasmids | Genotype | Source |
| *pLTetO1-dam* | *pLTetO1-dam ampR* | (Enrico Bena et al. 2024) |
| *pMVRedcas9_cIIg183a* | *pMVRed_cas9_cIIg183a_araC_neoR* | This study |
| *pLZ1* | *pBR322, bor::cm-24tetO* | (Trinh et al. 2020) |
| *pLZ3* | *pACYC177-pFtski-tetR-mCherry* | (Trinh et al. 2020) |
| pUC19 | *lacZα ampR* | (Norrander et al. 1983) |

| lambda phages | Genotype | Source |
|---|---|---|
| WT | Lambda *Ind1* $cI_{857}$ | Lab collection |
| Lambda tetO | Lambda $cI_{857}$ *bor::cat 24x tetO* | This study |
| lambda lysates | Genotype | Source |
| WT_L1 | Lambda *Ind1* $cI_{857}$ | Induction in MD114 |
| WT_L2 | Lambda *Ind1* $cI_{857}$ | Induction in MV235 |
| G183A_L1 | Lambda *Ind1* $cI_{857}$ $cII_{G183A}$ | Induction in MV318 |
| G183A_L2 | Lambda *Ind1* $cI_{857}$ $cII_{G183A}$ | Induction in MV319 |

**Table S9: List of oligos used for PCR, Gibson assembly and Sanger sequencing**

| Primer Name | Sequence 5'-3' | Source |
|---|---|---|
| cII_up | ACCACACCTATGGTGTATGCA | This study |
| cII_dwn | GTCATAATGACTCCTGTTGA | This study |
| JThM_50 | CTACAGTTATGGCGGAAAG | This study |
| JThM_51 | TATCAAGCAGCAGAATCATC | This study |
| JThM_52 | GTTTGATCAGAAGGACGTTG | This study |
| QF | GAGTGCGGAAGATGCAAAGG | (Fogg et al. 2010) |
| QR | TTAACAGTGCGTGACCAGG | (Fogg et al. 2010) |
| gyrB_F | GTCGAAGTGGCGTTGCAGTG | (Fogg et al. 2010) |
| gyrB_R | AGCCTGCCAGGTGAGTACCG | (Fogg et al. 2010) |
| stfR_up | GTGGGTGTCGGTTATAAG | This study |
| stfR_dwn | TATGGTCCGTGGTTGTTC | This study |
| tfaQ_up | TCCTCCTTAGTTCCTATTCC | (De Paepe et al. 2014) |
| tfaQ_dwn | CTGCCACTCATCGCAGTAC | (De Paepe et al. 2014) |
| polB_up | GATGGCAAAGCATTCGTCAC | This study |
| polB_dwn | ATGGCGCGAAGGCATATTAC | This study |
| dinB_up | GTGTTGAGAGGTGAGCAATG | This study |
| dinB_dwn | TAGAAAGAACCGCAACCG | This study |
| umuDC_up | CCACGTGAGCACAAGATAAG | This study |
| umuDC_dwn | GTTCGGTCGCTAATCCATTC | This study |
| dam_up | ACCTCGGCGCAATTTGTTTC | This study |
| dam_dwn | TCCTGGTTTCTGGCGTGTAC | This study |
| OMV310 | GCTTGCTGTTCTTGAATGGGGTTTTAGAGCTAGAAATAGCAAGTTAAAA | This study |
| OMV312 | GTTGTTCCATGCACCACAGGCCCATGGA | This study |
| OMV278 | ATGGAACAACGCATAACCCTGAAAG | This study |
| OMV279 | CGTCAACGACTCCCCATTCAAGAAC | This study |
| OMV280 | CTTGAATGGGGAGTCGTTGACGACG | This study |
| OMV281 | GAGGGATGCACCATTCTGAGATGTTTTT | This study |
| OMV308 | cctgtggtgcATGGAACAACGCATAACCC | This study |
| OMV309 | ccattcagcaGAGGGATGCACCATTCTG | This study |
| OMV304 | tgcatccctcTGCTGAATGGAAGCTTGG | This study |
| OMV314 | aacagcaagcGCTAAGATCTGACTCCATAAC | This study |
| OMV311 | agatcttagcGCTTGCTGTTCTTGAATGGGG | This study |

**Table S10. Rate of YFP-MutL foci**

The imaging time interval was 2 min (up) or 1 min (down). The "track length" is the time between the birth and cell division (or birth to lysis). The average track length was computed over all the considered cells +/- 2*SEM. μ: mean rate of YFP-MutL foci, $s_{\mu}$ = 2* SEM (Standard Error of the Mean). [1](Enrico Bena et al. 2024).

| Strain | Interval time (min) | Temperature | μ ± $s_{\mu}$ (/generation or infection) | N foci | N cell tracks | Mean track length (min) |
|---|---|---|---|---|---|---|
| 63ME120RL (6300 lambda) | 2 | 39 °C | (8.2 ± 0.6) | 2833 | 347 | (52.9 ± 1.2) |
| L12 (lambda *mutS*) | 2 | 39 °C | (0.04 ± 0.02) | 9 | 201 | (69.3 ± 1.5) |
| L6 (lambda homDlp12) | 2 | 39 °C | (9.1 ± 0.9) | 1233 | 136 | (64 ± 2) |
| 63ME120R (6300)[1] | 2 | 37 °C | (0.52 ± 0.004) | (313 ± 2) | 600 | (25.8 ± 0.03) |
| L6 (lambda homDlp12) | 2 | 35 °C | (0.37 ± 0.05) | 237 | 633 | (28.2 ± 0.4) |
| ME120RL (WT lambda) | 1 | 39 °C | (11.1 ± 0.9) | 3261 | 291 | (54.7 ± 0.7) |
| L6 (lambda homDlp12) | 1 | 39 °C | (12.4 ± 1.4) | 1787 | 144 | (63.7 ± 0.9) |
| 63ME120RL (6300 lambda) | 1 | 35 °C | (0.76 ± 0.18) | 108 | 143 | (30 ± 1) |

### References

Cox RS, Surette MG, Elowitz MB (2007) Programming gene expression with combinatorial promoters. Mol Syst Biol 3:. https://doi.org/10.1038/msb4100187

De Paepe M, Hutinet G, Son O, et al (2014) Temperate Phages Acquire DNA from Defective Prophages by Relaxed Homologous Recombination: The Role of Rad52-Like Recombinases. PLoS Genet 10:. https://doi.org/10.1371/journal.pgen.1004181

Dillon MM, Sung W, Lynch M, Cooper VS (2018) Periodic variation of mutation rates in bacterial genomes associated with replication timing. MBio 9:1–15. https://doi.org/10.1128/mBio.01371-18

Dillon MM, Sung W, Lynch M, Cooper VS (2015) The rate and molecular spectrum of spontaneous mutations in the GC-rich multichromosome genome of Burkholderia cenocepacia. Genetics 200:935–946. https://doi.org/10.1534/genetics.115.176834

Dillon MM, Sung W, Sebra R, et al (2017) Genome-wide biases in the rate and molecular spectrum of spontaneous mutations in vibrio cholerae and vibrio fischeri. Mol Biol Evol 34:93–109. https://doi.org/10.1093/molbev/msw224

Elez M, Radman M, Matic I (2012) Stoichiometry of MutS and MutL at unrepaired mismatches in vivo suggests a mechanism of repair. Nucleic Acids Res 40:3929–3938. https://doi.org/10.1093/nar/gkr1298

Enrico Bena C, Ollion J, De Paepe M, et al (2024) Real-time monitoring of replication errors' fate reveals the origin and dynamics of spontaneous mutations. Nat Commun 15:. https://doi.org/10.1038/s41467-024-46950-0

Fogg PCM, Allison HE, Saunders JR, McCarthy AJ (2010) Bacteriophage Lambda: a Paradigm Revisited. J Virol 84:6876–6879. https://doi.org/10.1128/jvi.02177-09

Foster PL, Hanson AJ, Lee H, et al (2013) On the mutational topology of the bacterial genome. G3 Genes, Genomes, Genet 3:399–407. https://doi.org/10.1534/g3.112.005355

Grant GN, Jessee J, Bloom FR, Hanahan D (1990) Differential plasmid rescue from transgenic mouse DNAs into Escherichia coli methylation-restriction mutants. Proc Natl Acad Sci U S A 87:4645–4649. https://doi.org/10.1073/pnas.87.12.4645

Herman C, Ogura T, Tomoyasu T, et al (1993) Cell growth and λ phage development controlled by the same essential Escherichia coli gene, ftsH/hflB. Proc Natl Acad Sci U S A 90:10861–10865. https://doi.org/10.1073/pnas.90.22.10861

Long H, Sung W, Miller SF, et al (2014) Mutation rate, spectrum, topology, and context-dependency in the DNA mismatch repair-deficient Pseudomonas fluorescens ATCC948. Genome Biol Evol 7:262–271. https://doi.org/10.1093/gbe/evu284

Martinsohn JT, Radman M, Petit MA (2008) The λ red proteins promote efficient recombination between diverged sequences: Implications for bacteriophage genome mosaicism. PLoS Genet 4:. https://doi.org/10.1371/journal.pgen.1000065

Niccum BA, Lee H, MohammedIsmail W, et al (2018) The spectrum of replication errors in the absence of error correction assayed across the whole genome of escherichia coli. Genetics 209:1043–1054. https://doi.org/10.1534/genetics.117.300515

Norrander J, Kempe T, Messing J (1983) Construction of improved M13 vectors using oligodeoxynucleotide-directed mutagenesis. Gene 26:101–106. https://doi.org/10.1016/0378-1119(83)90040-9

Rajagopala S V., Casjens S, Uetz P (2011) The protein interaction map of bacteriophage lambda. BMC Microbiol 11:213. https://doi.org/10.1186/1471-2180-11-213

Robert L, Ollion J, Robert J, et al (2018) Mutation dynamics and fitness effects followed in single cells. Science (80- ) 359:1283–1286. https://doi.org/10.1126/science.aan0797

Schaaper RM, Dunn RL (1987) Spectra of spontaneous mutations in Escherichia coli strains defective in mismatch correction: the nature of in vivo DNA replication errors. Proc Natl Acad Sci U S A 84:6220–6224. https://doi.org/10.1073/pnas.84.17.6220

Sergueev K, Court D, Reaves L, Austin S (2002) E. coli cell-cycle regulation by bacteriophage lambda. J Mol Biol 324:297–307. https://doi.org/10.1016/S0022-2836(02)01037-9

Shao Y, Wang IN (2008) Bacteriophage adsorption rate and optimal lysis time. Genetics 180:471–482. https://doi.org/10.1534/genetics.108.090100

Taylor K, Węgrzyn G (1995) Replication of coliphage lambda DNA. FEMS Microbiol Rev 17:109–119. https://doi.org/10.1016/0168-6445(95)00077-1

Trinh JT, Shao Q, Guan J, Zeng L (2020) Emerging heterogeneous compartments by viruses in single bacterial cells. Nat Commun 11:1–11. https://doi.org/10.1038/s41467-020-17515-8

Wold MS, Mallory JB, Roberts JD, et al (1982) Initiation of bacteriophage λ DNA replication in vitro with purified λ replication proteins. Proc Natl Acad Sci U S A 79:6176–6180. https://doi.org/10.1073/pnas.79.20.6176

Zaritsky A, Wang P, Vischer NOE (2011) Instructive simulation of the bacterial cell division cycle. Microbiology 157:1876–1885. https://doi.org/10.1099/mic.0.049403-0
